# Ultra-weak protein-protein interactions can modulate proteome-wide searching and binding

**DOI:** 10.1101/2022.09.30.510365

**Authors:** Jennifer L. Hofmann, Akshay J. Maheshwari, Alp M. Sunol, Drew Endy, Roseanna N. Zia

## Abstract

Research on protein-protein interaction (PPIs) tends to focus on high affinity interactions. Weaker interactions (*K*_*d*_ >1*μM*) recently understood as contributing to intracellular phase separation suggest that even-weaker PPIs might also matter in as-yet unknown ways. However, ultra-weak PPIs (*K*_*d*_ >1*mM*) are not readily accessible by *in vivo* techniques. Here we use protein electrostatics to estimate PPI strengths and spatially-resolved dynamic simulations to investigate the potential impacts of ultra-weak PPIs within dense protein suspensions. We find that ultra-weak PPIs can drive formation of transient clusters that last long enough to enable enzyme-catalyzed reactions and accelerate the sampling of protein associations. We apply our method to *Mycoplasma genitalium*, finding that ultra-weak PPIs should be ubiquitous among cytoplasmic proteins. We also predict that the proteome-wide interactome can be shifted to favor ‘binding-dominant’ ultra-weak PPIs via the introduction of a few charged protein complexes. We speculate that ultra-weak PPIs could contribute to cellular fitness by facilitating sampling and colloidal-scale transport of proteins involved in biological processes, including protein synthesis.

## Introduction

Protein-protein interactions (PPIs) are essential for life, allowing molecules to interact and function together. Multi-protein complexes, including ribosomes, polymerases, and cytoskeletal filaments, organize the crowded cellular interior and form complex biomolecular networks for carrying out cellular functions. These networks of interactions are distributed physically throughout the cytoplasm and regulate the most fundamental of biological processes, from control of gene expression (Grüner et al., 2016) to cell signaling, growth, and oncogenesis (Tran et al., 2021).

The study of PPIs has traditionally focused on interactions that correlate with a strong observable phenotype and thus are often high affinity interactions (*K*_*d*_ < 1*μM*). The advent of high-throughput methods, most notably yeast two-hybrid (Y2H) (Brückner et al., 2009; Trigg et al., 2017), protein-fragment complementation (PCA) (Schlecht et al., 2017; Liu et al., 2020), proximity labeling (PL) (Qin et al., 2021), and mass spectrometry-based (Low et al., 2021; Smits and Vermeulen, 2016) assays, has enabled proteome-wide screening of thousands to millions of interactions. Such methods primarily focus on PPI detection but can be expanded or paired with biophysical techniques, such as fluorescence correlation spectroscopy (FCS), Förster resonance energy transfer (FRET), and nuclear magnetic resonance (NMR) spectroscopy, to more precisely quantify the strength of any given interaction. Improvements in these methods have enabled detection of ever-weaker interactions and their connections to cellular function. At the current limit of what can be readily observed *in vivo*, weak interactions (*K*_*d*_ > 1*μM*, **Figure 1**) have been found to underlie intracellular phase separation and formation of membraneless organelles, impacting biological processes including gene expression, signaling, and stress response (Brangwynne et al., 2009; Feric et al., 2016; Li et al., 2012; Lyon et al., 2020).

**Figure 1.**
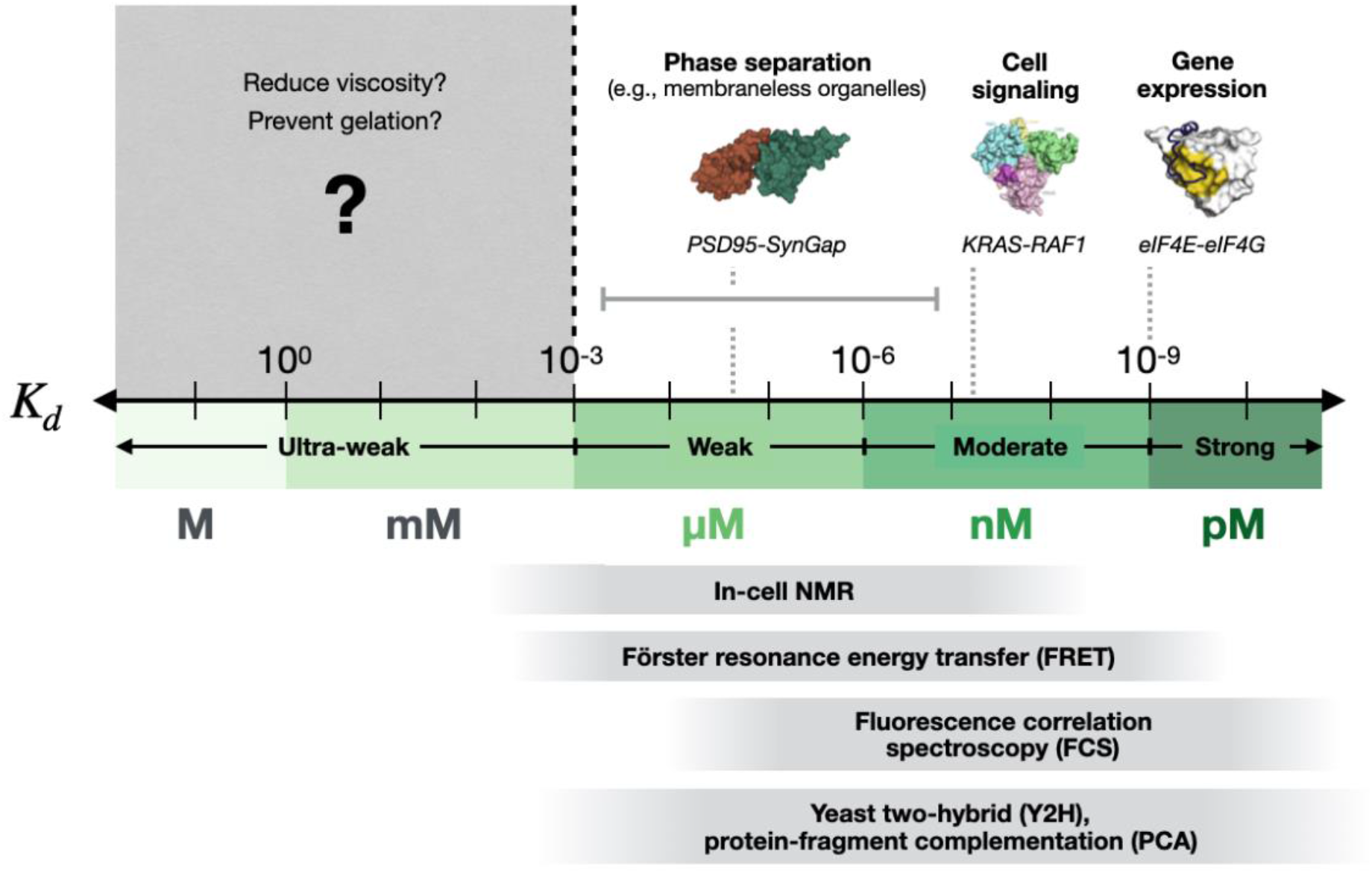
Protein-protein interactions are important across all measurable affinities. Might ultra-weak interactions that cannot now be readily measured matter, too? The binding affinity of proteins is quantified via the equilibrium dissociation constant, *K*_*D*_. Experimental techniques used to quantify binding affinity can readily measure strong and moderate interactions (accessible ranges, light grey, bottom) (Durfee et al., 1993; Lin et al., 2018; Mustafi and Weisshaar, 2018; Sugiki et al., 2019), but limited temporal and spatial resolution largely prevents *in vivo* study of interactions in the ultra-weak millimolar range (dark grey, top). *K*_*d*_ and reference protein structures are adapted with permission from (Grüner et al., 2016; Li et al., 2012; Tran et al., 2021; Zeng et al., 2016), including interactions between postsynaptic density protein 95 (PSD-95) and SynGAP which underlie phase separation in neurons (Zeng et al., 2016), between two signaling proteins KRAS and RAF1 involved in oncogenesis (Tran et al., 2021), and between two eukaryotic initiation factors 4E and 4G (eIF4E-eIF4G) involved in translation (Grüner et al., 2016).

Such observations and discoveries suggest that understanding even-weaker PPIs might reveal further contributions to cellular function. However, existing *in vivo* techniques are largely limited to detecting interactions with sub-millimolar affinities. Work to increase the sensitivity of high-throughput methods such as Y2H and PCA to include weak, transient PPIs can result in false positives (Brückner et al., 2009). Moreover, inherent physico-chemical constraints can bias assays towards detection of certain types of PPIs (e.g., those involving transmembrane or nuclear proteins), leading to high false negative rates (Jensen and Bork, 2008). As a result, such methods can most reliably be used to discover interactions with micromolar or stronger affinities and must be carefully chosen to target desired PPIs (Durfee et al., 1993). Three common biophysical techniques for quantifying PPIs *in vivo* – FCS, FRET, and NMR-based methods – also require introduction of perturbations that can interfere with weak PPIs (Chien and Gierasch, 2014) and are inherently limited in their spatial and temporal resolution (Algar et al., 2019; Freedberg and Selenko, 2014; Lin et al., 2018; Mustafi and Weisshaar, 2018; Sugiki et al., 2019; Sukenik et al., 2017) (**Figure 1**). Thus, ‘ultra-weak’ PPIs, with millimolar *K*_*d*_, represent an under-studied interaction regime inside cells.

At first glance, a biologist might expect such ultra-weak PPIs (UW-PPIs) do not last long enough to matter (i.e., impact cell phenotype or fitness). However, from a first-principles physics perspective it is well known that ultra-weak interactions among molecules in suspension can impact the performance of the ensemble. For example, in inorganic systems, interactions that are close in strength to the energy of spontaneous Brownian fluctuations are strong enough to ‘pull’ particles through a suspension and can reduce overall viscosity, accelerate sedimentation, and prevent gelation (Hoh and Zia, 2016b, 2016a; Huang and Zia, 2019; Moncho-Jordá et al., 2010; Stradner et al., 2007). Moreover, the so-called “quinary structure” of proteins (McConkey, 1982), induced by the complex intracellular milieu, has been demonstrated to impact protein folding and stability relative to dilute buffer (Feng et al., 2019; Monteith et al., 2015). Recent *in vitro* studies using advanced centrifugation- (Rowe, 2011) and NMR-based (Johansson et al., 2014; Milles et al., 2015; Xing et al., 2014) methods have also quantified a handful of individual PPIs with very low affinities, including an interaction between nucleoprotein & phosphoprotein (*K*_*d*_ ∼0.6 *mM*), which is essential for measles virus replication (Milles et al., 2018), and between adaptor proteins NcK-2 and PINCH-1 (*K*_*d*_ ∼3 *mM*), which ensures proper eukaryotic cell adhesion (Vaynberg et al., 2005). These findings suggest that UW-PPIs, which are now largely inaccessible by *in vivo* methods, could have broader impacts on the emergent behavior of cellular systems.

Here, we adapt modeling and simulation tools that are well-established for inorganic colloidal systems to more formally explore the extent to which we would expect UW-PPIs to impact cellular behavior. Colloidal systems are comprised of “colloids,” which are nanometer- to micron-sized macromolecules that undergo Brownian diffusion when suspended in a liquid. Specifically, we use Brownian dynamics simulations to interrogate the organizational and dynamic impacts of UW-PPIs within dense cell-like environments. Our results show that UW-PPIs can drive transient protein co-localization while permitting ongoing protein diffusion, enabling more rapid sampling of protein associations with durations on the order of enzyme-catalyzed reactions. We extend our results to the cytoplasmic proteome of *Mycoplasma genitalium*, a model prokaryote for minimal genome research, and find a prevalence of ‘binding-dominant’ UW-PPIs, particularly relevant for proteins involved in tRNA synthesis, metabolism, and protein folding. We hypothesize that interactions in this binding-dominant UW-PPI regime might undergird combinatoric sampling within cytoplasm that could be essential for optimal growth and fitness.

## Results

### I. Computational modeling of ultra-weak PPIs

Protein-protein interactions (PPIs) are most often reported as bulk binding affinities that quantify the strength of an interaction in terms of how easily a complex dissociates. Such equilibrium dissociation constants, *K*_*D*_, are defined kinetically as the ratio of the dissociation and association rate constants, or thermodynamically as a function of the relative concentrations of bound and free proteins (**Figure 1**). However, the strength of a PPI can also be understood in terms of its mechanistic origin: a dynamic competition between deterministic attractions that tend to pull proteins together (*V*_0_, the attraction strength at protein-protein contact), versus the entropic, stochastic thermal fluctuations underlying Brownian motion that tend to drive proteins apart (*k*_*B*_*T*, where *k*_*B*_ is the Boltzmann constant and *T* is the absolute temperature) (**Figure 2A**). In the case of UW-PPIs, these two energies are similar in strength and thus such interactions are transient.

**Figure 2.**
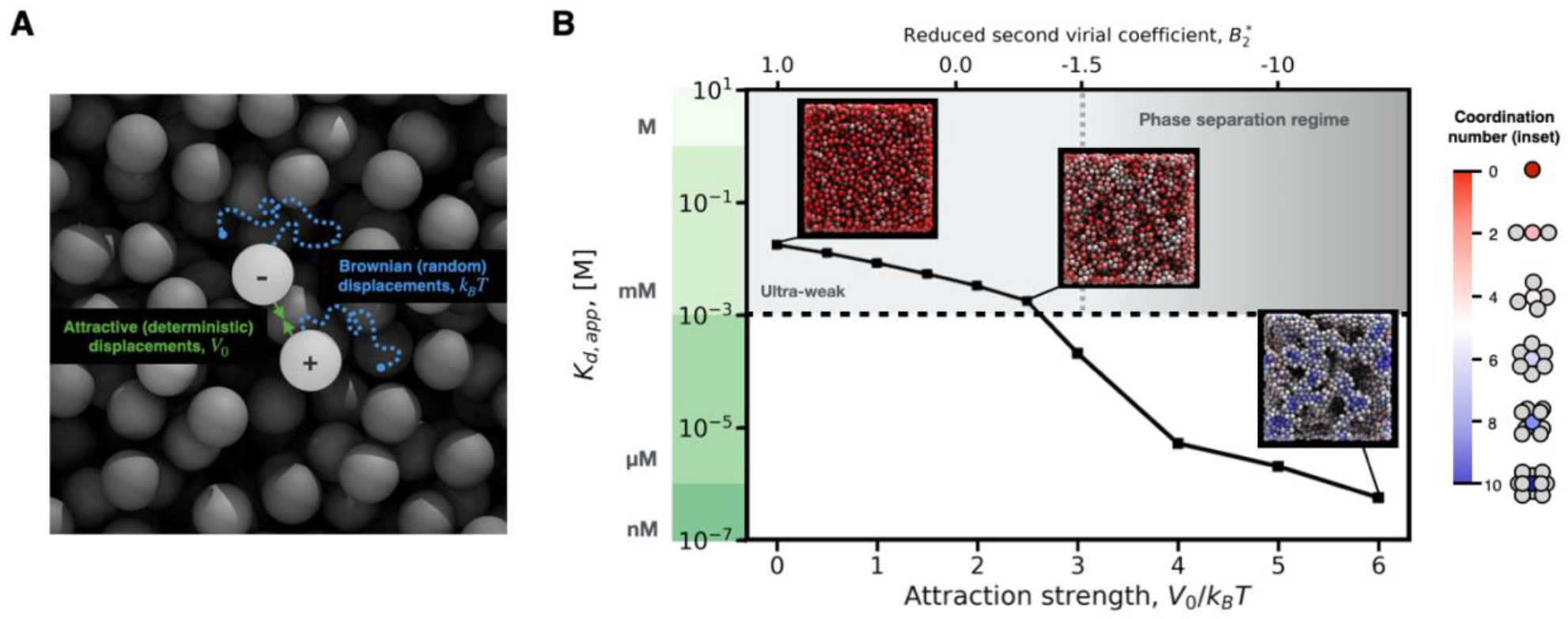
Ultra-weak protein-protein interactions can produce low-order multi-protein clusters. **(A)** A schematic representation of the competition between attractions, characterized by the strength at contact *V*_0_, and the thermal energy, *k*_*B*_*T*, that drives Brownian motion of colloidal-scale particles in a suspension. The transient nature of UW-PPIs, which are close in magnitude to the underlying thermal energy, emerges from this competition. **(B)** An increase in the interparticle attraction strength relative to thermal energy, corresponds with a stronger apparent binding affinity, *K*_*d,app*_, with values including the ultra-weak regime (shaded grey region, above dashed black line). Liquid-liquid phase separation can occur for systems with a reduced second virial coefficient, 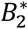, less than −1.5 (Noro and Frenkel, 2000; Vliegenthart and Lekkerkerker, 2000) (x-axis, upper; dark grey shaded region to right of vertical dashed line). Simulation snapshots (inset, colored) highlight how proteins are coordinated with more neighbors as attraction strength increases, forming transient clusters and durable gel networks.

As a representative biological system, we chose to study the cytoplasmic proteome of *M. genitalium*, a model organism for minimal genome research and whole-cell computational modeling (Gibson et al., 2008; Karr et al., 2012). Though there are different physical origins of PPIs, we focused on electrostatic attractions because they are ubiquitous across charged proteins, are known drivers of cellular processes, and manifest over a range of strengths *in vivo* (Mihiri Shashikala et al., 2019; Pabbathi et al., 2022; Schavemaker et al., 2017). Using estimates of protein charges as determined by all-atom simulation (Dutagaci et al., 2021; Feig et al., 2015), we calculated the strengths of all pair-wise electrostatic attractions between cytoplasmic proteins and durable protein-RNA complexes (e.g. ribosomes) in *M. genitalium* (see **Methods**). We found these predicted strengths span a dynamic range from zero to six times the strength of thermal fluctuations (i.e., *V*_0_/*k*_*B*_*T* = 0 to 6).

We simulated the physical dynamics of protein-protein interactions in a dense suspension representative of prokaryotic cytoplasm at moderate growth rate (i.e., 30% occupied volume fraction, see **Methods**), with thousands of proteins modeled individually as physical objects diffusing in the cytosolic liquid. In our model, the microscopic diffusion of individual proteins emerges from a random-walk process arising from explicitly represented stochastic displacements obeying Gaussian statistics, rather than an imposed bulk diffusion coefficient (see **Movie S1, Methods**). Entropic exclusion arising from the discrete size of each protein hinders diffusion proportionally to the overall occupied volume fraction of the system. We also endowed each protein with an electrostatic surface potential that produces a deterministic attraction of strength zero to six times the strength of thermal fluctuations (see dynamic range above). Throughout each simulation, the positions of individual proteins were then tracked as they diffuse, bind, and unbind within the crowded milieu, experiencing the interplay between physico-chemical attractions and Brownian fluctuations that underlie weak PPI associations (**Figure 2A**).

### II. How are physical forces related to dissociation constants?

We next determined a quantitative mapping of the mechanistic physico-chemical attraction strength relative to thermal energy, *V*_0_/*k*_*B*_*T*, onto the apparent equilibrium dissociation constant, *K*_*d,app*_, the using the computed concentrations of bound proteins (**Figure 2B, Methods**). We demonstrated that dynamic colloidal simulations can be used to estimate expected protein-protein affinities across a dynamic range of interaction strengths. Importantly, we showed that interactions in the now experimentally inaccessible ultra-weak regime can be computed (**Figure 2B**, above dashed horizontal line); specifically, we showed the physico-chemical threshold for UW-PPIs to be about three times the strength of thermal fluctuations (*V*_0_/*k*_*B*_*T* = 2.7).

We then looked at even weaker interactions, specifically at the limit of no attraction (i.e., *V*_0_/*k*_*B*_*T* = 0, **Figure 2B**). One might expect an infinitely large dissociation constant, but our simulations revealed a measurable apparent affinity of *K*_*d,app*_ ∼ 20 mM. This finite affinity arises solely from the non-zero likelihood that objects exerting no deterministic attractions will diffuse near each other for short time intervals. Eventually, a protein pair will drift apart, as the stochastic Brownian motion acts to make uniform their spatial distribution (Batchelor, 1976). Even in the absence of deterministic attractions, these entropic protein-protein ‘encounters’ produce apparent interactions that can be measured.

### III. Microstructural effects of ultra-weak PPIs

Entropic exclusion affects the spatial arrangement of proteins (known in the colloidal physics literature as the suspension ‘microstructure’ (Morris, 2009; Russel et al., 1989)). This phenomenon is easily seen in the suspension radial distribution function, where the most likely position of a protein near a tracer is in the shell of nearest-neighbors – even without any attractive forces (**Figure S1B**). This short-range co-localization frees up the larger surrounding volume to maximize long-range entropy, while also ensuring that a pair of proteins close to each other will soon separate to sample the rest of the configuration space. Even at the zero-attraction limit (*V*_0_/*k*_*B*_*T* = 0), we found that at least 22% of proteins are members of transient dimers and trimers (**Figure S3)**.

At the opposite limit, in the moderate-affinity case (*V*_0_/*k*_*B*_*T* = 6), proteins are bonded to many other neighbors, forming a system-spanning proteinaceous gel. This co-localization is visible in snapshots from dynamic simulations (insets, **Figure 2B**); each spherical protein is colored according to the number of other proteins with which it is interacting (i.e., coordinated) at a given instant (color bar, right). In between the moderate-affinity and zero-attraction limits, we found that PPIs at the ultra-weak threshold (*V*_0_/*k*_*B*_*T* = 2.5) can also produce measurable complexation. The images show formation of higher-order clusters, reducing the total fraction of monomers from 83% in the zero-attraction case to 53% at the ultra-weak affinity threshold (**Figure S3B**). Overall, our results suggest that UW-PPIs operate at a ‘sweet spot’ of stability, allowing co-localization but not locking in long-range network structure.

We next asked how UW-PPIs compare in strength to the interactions known to underlie intracellular phase separation and formation of membraneless organelles (Brangwynne et al., 2009; Feric et al., 2016; Li et al., 2012; Lyon et al., 2020). To do so, we calculated the reduced second virial coefficient 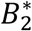 (see **Methods**), which predicts the onset of phase separation in colloidal suspensions. 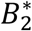 is a convenient scalar that quantifies the overall balance of attraction and repulsion in an interaction, independent of its physico-chemical origin (e.g., electrostatics, hydrophobicity) (Noro and Frenkel, 2000). When 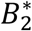 is zero, proteins are effectively ‘point particles’ (i.e. an ideal gas) and do not interact at all. When 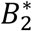 reaches approximately −1.5 (Noro and Frenkel, 2000; Vliegenthart and Lekkerkerker, 2000), attractions cause liquid-liquid phase separation (**Figure 2B**, upper x-axis). In our simulations, we found that in the hard-sphere zero-attraction case, 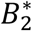 is positive. Entropic exclusion leads to an effectively ‘repulsive’ regime that assures particles do not overlap (i.e., have finite-rather than point-size). As affinity increases, the suspension becomes more ‘attractive,’ reducing 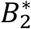. We found that UW-PPIs correspond with 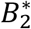 greater than −0.7 (**Figure 2B**, upper x-axis), and thus drive transient cluster formation rather than full liquid-liquid phase separation. Moreover, these 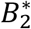 correspond to the values shown previously to reduce overall viscosity in inorganic suspensions (Huang and Zia, 2019), suggesting that UW-PPIs are sufficiently strong to ‘pull’ proteins through the cellular milieu (Huang et al., 2022).

### IV. Dynamic effects of ultra-weak PPIs

To better understand if UW-PPIs can last long enough to matter for biological functions, we estimated the time proteins spend interacting with one another (‘binding events’) relative to searching for new partners (‘search events’). To do so, we tracked individual separation trajectories of protein pairs and plotted the trends in binding and search event durations for three cases: (i) hard-sphere zero-attraction, (ii) ultra-weak threshold, and (iii) moderate-affinity (**Figure 3A**, upper; see **Methods** for calculation). The result is an explicit dynamical prediction of interactions, without the simplifying ‘well-mixed’ and ‘point-particle’ assumptions common in the literature (Andrews and Bray, 2004; Grima and Schnell, 2006; Smoluchowski, 1916). In addition, we quantified the lifetime of trimers and larger clusters by tracking the average cluster dissociation time, *τ*_*cluster*_, in each system (**Figure 3B, S7B**; see **Methods**).

**Figure 3.**
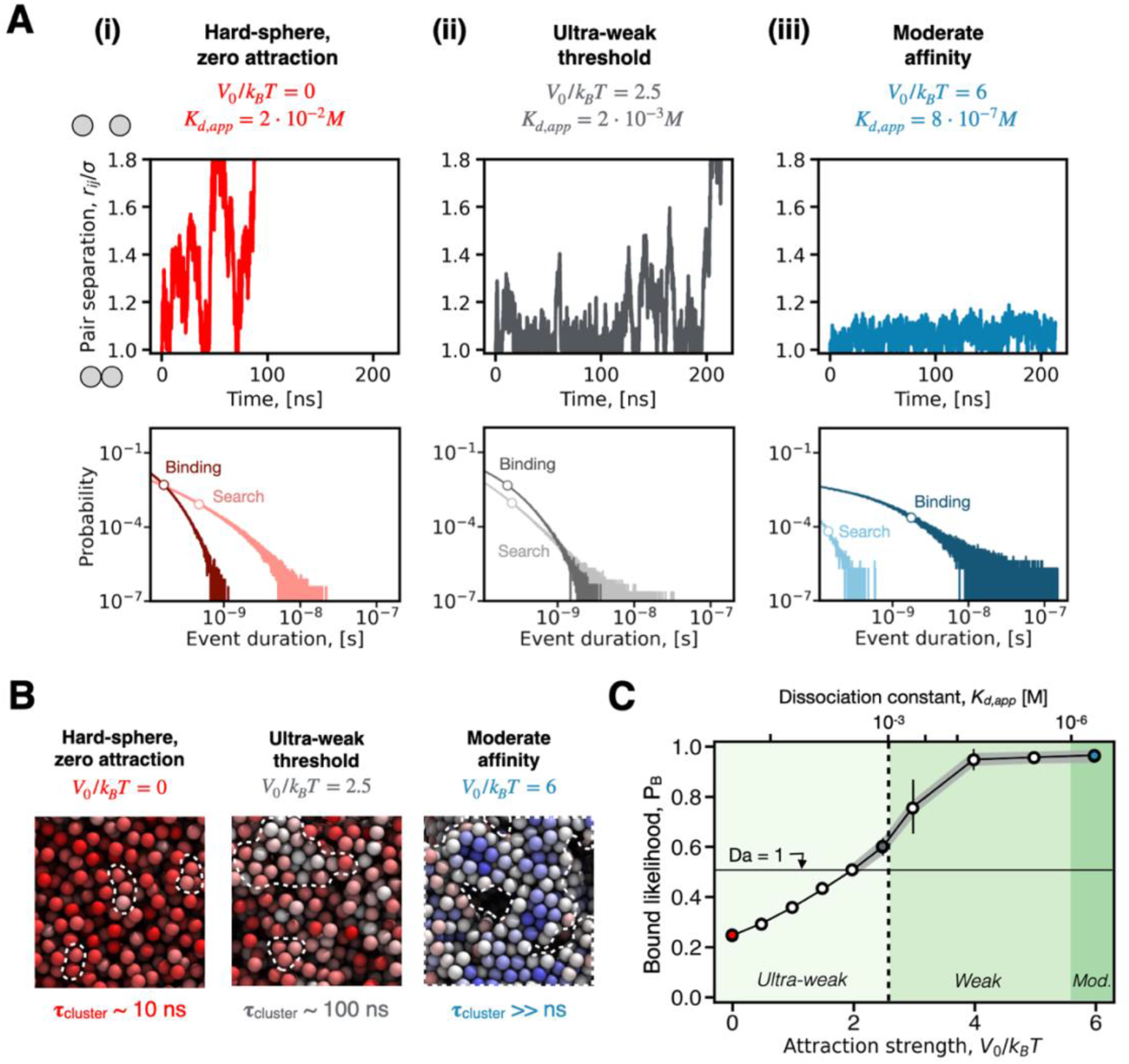
Ultra-weak protein-protein interactions can transition the cytoplasmic experience of individual proteins from a search-dominant to binding-dominant regime. **(A) (Upper)** Example separation trajectories for individual protein pairs, as a function of time. Pairs begin bound (close to contact i.e., center-to-center separation distance *r*_*ij*_/*σ* = 1, where *σ* is the protein diameter) and their separation distances are tracked as they bind and unbind. Protein pairs separate rapidly in the hard-sphere, zero-attraction case (red) compared with the ultra-weak (grey) and moderate affinity (blue) cases. **(Lower)** Ensemble probability distributions for the time spent by proteins bound to one another (‘binding’ events, dark plots) and searching for new partners (‘search’ events, light plots). As binding affinity increases, left to right, associations are longer-lasting and search events are more rapid **(B)** Proteins form transient clusters and durable networks (white dashes) as affinity increases, with the average cluster dissociation time shown for each affinity. A PPI at the ultra-weak affinity threshold (grey) drives clusters that last an order of magnitude longer than the hard-sphere case (red) but are still dynamic compared to the moderate affinity case (blue). **(C)** As affinity increases (inset colors, light to dark), proteins are more likely on average to be found in a bound state (‘bound likelihood’ *P*_*B*_). A transition from a ‘search-dominant’ to a ‘binding-dominant’ regime occurs when proteins spend more than half of their time in encounters (Damköhler number Da = 1, grey highlighted section above horizontal line). This transition occurs within the ultra-weak regime (left of the vertical dashed line). Red, grey, and blue points correspond to conditions in (B).

We found that, even without attractions, proteins experience many fleeting, yet measurable encounters driven only by entropy and Brownian motion (**Figure 3A(i)**). In this zero-attraction limit, non-attractive ‘binding’ events are finite but shorter than search events (**Figure 3A(i)**, lower, circular markers). Mechanistically, a pair of proteins is brought together by Brownian motion but quickly separate due to entropy unless attractive forces hold them together, allowing each protein to rapidly sample novel encounters with other proteins. The transient nature of weakly bound encounters also holds at the level of interaction networks (i.e., the mesoscale). For example, in the hard-sphere zero-attraction limit, we found that small clusters exist even though they dissociate rapidly, on the order of 10 ns (**Figure 3B**, left).

In the moderate-affinity limit (**Figure 3A(iii)**), the opposite is true: we found that proteins spend much more time bound together than diffusively searching alone. In this regime, it is easy to encounter a nearby protein owing to proximity in the surrounding condensate, but since the interaction strength is much stronger than Brownian motion, proteins rarely break free from each other and thus have fewer novel binding partners. Macroscopic condensates can thus last on the order of hours (Padmanabhan and Zia, 2018) and prevent large-scale restructuring of the percolated suspension (**Figure 3B**, right).

At the ultra-weak threshold (**Fig3A(ii)**), the interplay between co-localization and spontaneous Brownian fluctuations brings the search and binding times to nearly the same value, comparable to the timescales of elementary chemical reactions (Gruebele and Zewail, 1990) and protein folding (Bredenbeck et al., 2003). We inferred that UW-PPIs permit many subsequent associations following each binding event by increasing mesoscale proximity, noting that higher-order microstructure (clusters) can keep proteins closer over much longer timescales. We found that PPIs at the ultra-weak threshold (*V*_0_/*k*_*B*_*T* = 2.5) drive clusters that last an order of magnitude longer than in the hard-sphere, zero-attraction limit, with dissociation times on the order of 100 ns (**Figure 3B**, center). Such ultra-weak clusters are sufficiently durable to facilitate the fastest known enzyme-catalyzed reactions (Smejkal and Kakumanu, 2019) (please see Discussion, **Figure 5**). However, co-localization is still transient enough to maintain molecular mobility and enable the suspension to reorganize, unlike in the moderate-affinity case. Overall, UW-PPIs operate at a dynamical ‘sweet spot’ in between the hard-sphere limit, which allows for copious sampling but little time for chemistry to occur, and the moderate-affinity limit, where protein experience durable associations with limited sampling.

**Figure 5.**
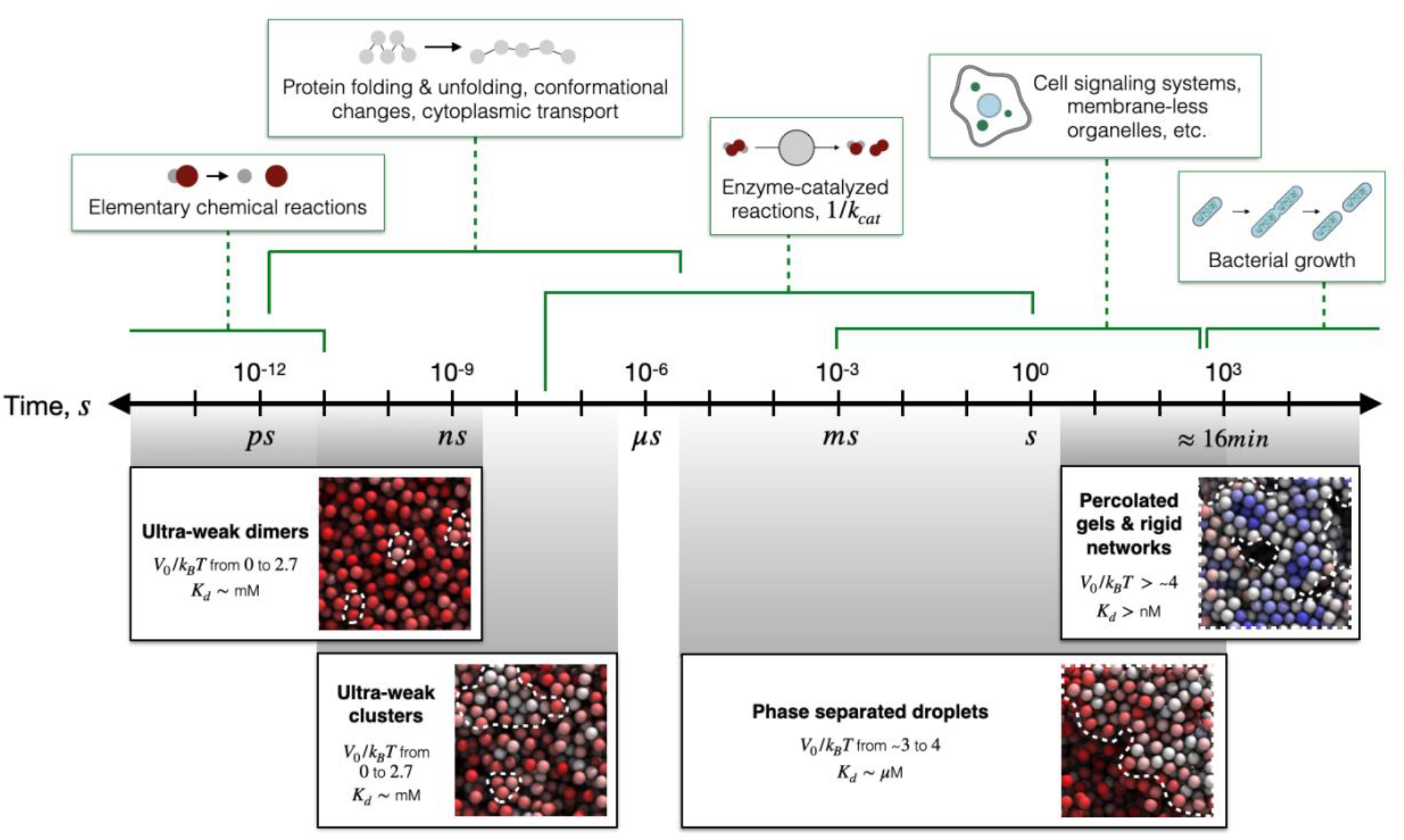
Ultra-weak PPIs below current detection limits can produce transient associations that are long-lived enough for enzyme-catalyzed reactions but are faster than other established biological phenomena. Intracellular phenomena occur over a range of timescales (upper, green), from bacterial division times as fast as 10 minutes (Rocha, 2004) down to physico-chemical fluctuations on the order of femto- to nano-seconds. The residence times of ultra-weak protein dimers (lower, left) are comparable to the timescales of elementary chemical reactions (Gruebele and Zewail, 1990) and protein conformational fluctuations (Bredenbeck et al., 2003), whereas higher-order clusters can last an order of magnitude longer (lower, center), on par with enzyme-catalyzed reactions (Smejkal and Kakumanu, 2019). This ultra-weak clustering regime operates below the timescales of phase separated droplets (Molliex et al., 2015; Tang et al., 2021), cell signaling (Asthagiri and Lauffenburger, 2003), and percolated networks (Ramm et al., 2022; Zia et al., 2014) (lower, right), which are sufficiently durable to be measured *in vivo*.

We next sought to better understand the bulk experience of proteins in our model and explore how PPIs can shift collective behavior of proteins in cytoplasm. To do so, we quantified the overall likelihood of finding a protein in a binding event (‘bound likelihood’, **Figure 3C**) to understand the average state of a protein at a given binding affinity. We found that entropic exclusion alone (*V*_0_/*k*_*B*_*T* = 0) causes proteins to spend 22% of their time bound, despite rapidly sampling many different partners (**Figure 3A**). As affinity increases, the overall bound likelihood reaches 61% at the ultra-weak limit (*V*_0_/*k*_*B*_*T* = 2.5). Eventually, at moderate affinity, proteins are nearly always bound to other proteins. Our results thus suggest that, despite their brevity relative to higher-affinity attractions, ultra-weak binding events can be numerous enough to cause proteins to spend most of their time bound in dynamic associations with surrounding proteins.

We quantified this balance between dynamic associations and diffusive search events via a Damköhler number (*Da* = *τ*_*search*_ /*τ*_*bind*_, where *τ*_*search*_ and *τ*_*bind*_ are the average duration of search and binding events, respectively). When *Da* < 1, proteins spend more than half of their lifetimes bound to other proteins; we termed this the ‘binding-dominant’ PPI regime (**Figure 3C**, grey highlighted trend). As expected, moderate-affinity interactions fall within this ‘binding-dominant’ category. Surprisingly, we found that a subset of UW-PPIs also has *Da* <1 (grey highlighted region to left of dashed line). These interactions, generally not thought of as important relative to high-affinity PPIs, can in fact be strong enough to shift protein behavior to a binding-dominant regime (*V*_0_/*k*_*B*_*T* = 2 to 2.7, **Figure 3C**).

### V. Ultra-weak binding-dominant PPIs in the *M. genitalium* cytoplasm

We next sought to understand the prevalence of UW-PPIs in a representative organism, returning to the cytoplasmic proteome of a naturally-occurring minimal cell, *M. genitalium*. As described in Section I, we calculated the strengths of all pair-wise electrostatic interactions in the cytoplasmic proteome using estimated charge data computed via all-atom models (Dutagaci et al., 2021) (see **Methods, Figure S10**). We noted that about half (42%) of all such pairings in the *M. genitalium* cytoplasm should involve proteins or complexes of opposite charges and thus be attractive in nature.

To start, we quantified the probability distribution of attraction strengths across pairs of proteins in the *M. genitalium* cytoplasm (**Figure 4A**, upper). We found that most isotropic electrostatic PPIs in *M. genitalium* are ultra-weak, representing 99% of all attractive pairings. Further, 2% of electrostatic attractions are ultra-weak yet are sufficiently strong to drive binding-dominant protein behavior (**Figure 4A**, between vertical solid and dashed lines). We also analyzed the distributions of electrostatic PPIs experienced for each individual cytoplasmic protein (**Figure 4A**, lower), finding that proteins involved in tRNA synthesis, metabolism, and protein folding are the most likely to experience binding-dominant UW-PPIs. PPIs with affinities in this range (**Table S1**) include interactions between two chaperone proteins involved with protein folding (GroEL/ES and DnaJ/DnaK/GrpE, with attraction strength *V*_0_/*k*_*B*_*T* = 2.35) and between pyruvate kinase and pyruvate dehydrogenase (with *V*_0_/*k*_*B*_*T* = 2.52) involved with glycolysis and entry into the citric acid cycle.

**Figure 4.**
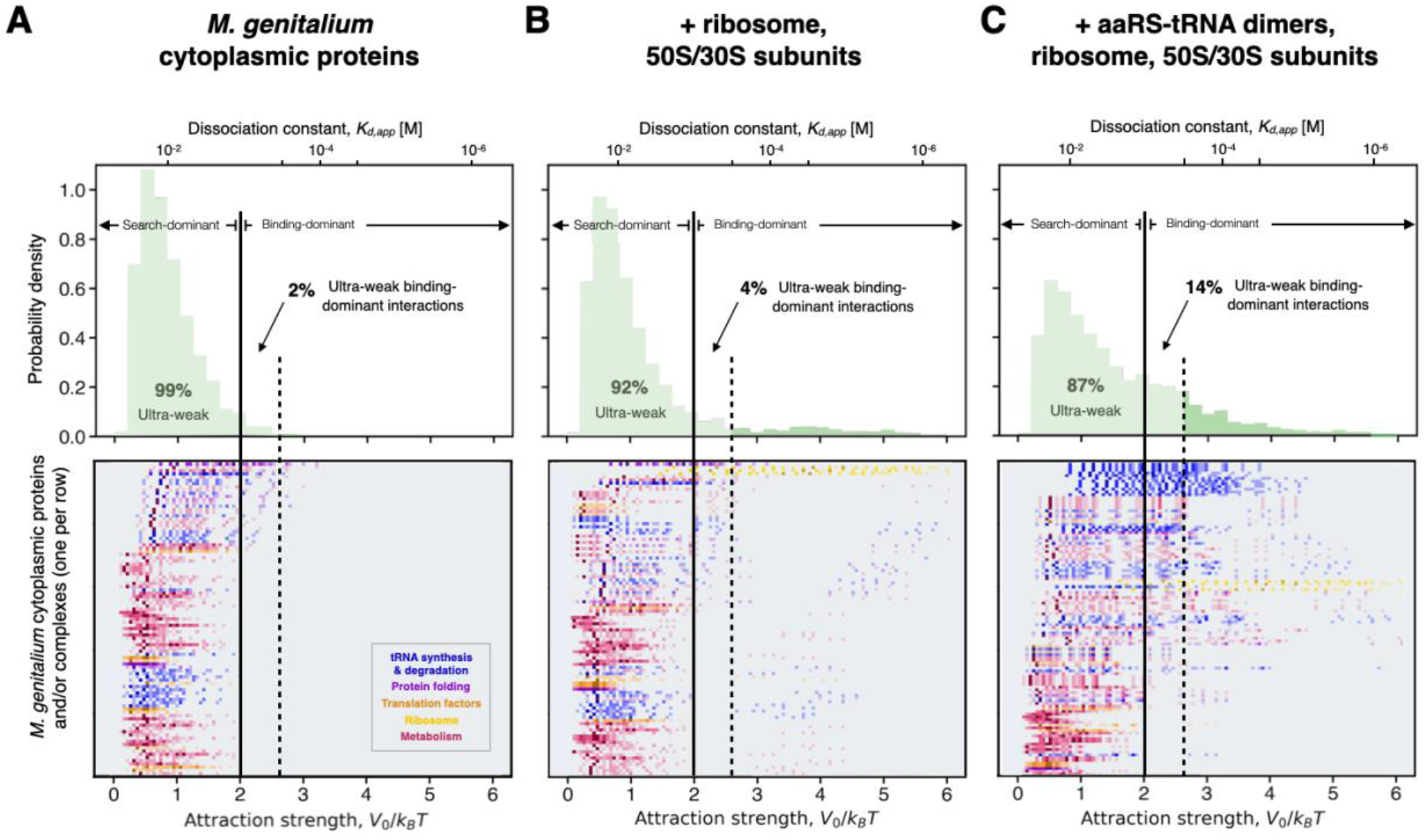
The vast majority of electrostatic attractive PPIs in the *M. genitalium* cytoplasm are ultra-weak and can be shifted into the binding-dominant regime by the addition of protein synthesis machinery. Probability distribution of pair-wise electrostatic attraction strengths between protein pairs in the *M. genitalium* cytoplasm **(A)** alone and **(B)** with the addition of ribosomes, 30S, & 50S subunits and **(C)** with the addition of ribosomes, 30S and 50S subunits, and aminoacyl-tRNA synthetase (aaRS) – tRNA dimers. (Upper) Interactions span a range of affinities, with the majority in each case classified as ultra-weak. A subset of interactions is strong enough to drive binding-dominant protein behavior (region between vertical solid and dashed lines), increasing from 2% to 14% of cytoplasmic interactions as more protein-RNA complexes are included. (Lower) Probability distribution of pair-wise electrostatic attraction strengths experienced by each protein or protein complex in the *M. genitalium* cytoplasm (where color intensity corresponds to probability, scaled as in the full upper distribution). Proteins in certain functional classes (colors: legend, inset) are more likely than others to experience binding-dominant UW-PPIs (rank-ordered, top to bottom).

Cytoplasmic proteins also interact with durable protein-RNA complexes, most notably the ribosome. We again calculated the probability distribution of attraction strengths, but further included interactions between cytoplasmic proteins and the 70S ribosome and its 30S and 50S subunits (**Figure 4B**, upper). We found that consideration of ribosomal complexes can shift the overall protein interaction distribution to higher attraction strengths, reducing the likelihood of UW-PPIs (92%) and doubling the likelihood of binding-dominant UW-PPIs (4%). Given their significant negative charge, ribosomal complexes experience stronger electrostatic interactions than most of the cytoplasmic proteome (**Figure 4B**, lower), including binding-dominant UW-PPIs with the ribosomal recycling factor, elongation factor 4, and initiation factor 1 (with *V*_0_/*k*_*B*_*T* = 2.34 to 2.65, see **Table S2**).

We next considered another set of important complexes: dimers between aminoacyl-tRNA synthetases (aaRSs), which are responsible for attaching the correct amino acid to tRNAs, and tRNAs. We found that the interaction distribution is again shifted to higher attraction strengths (**Figure 4C**, upper). Unlike the long tail which emerged from considering protein::ribosome interactions, protein::aaRS-tRNA interactions are primarily found in the binding-dominant ultra-weak regime (14%; **Figure 4C**, lower). We also noted that aaRS-tRNA complexes are likely to experience binding-dominant UW-PPIs with other aaRS that have not yet bound to tRNA (**Table S3**).

To compare the results of our electrostatics-based PPI prediction to experimental results, we examined the 13% of isotropic electrostatic attractions that are predicted to be strong enough to be detected by current *in vivo* methods (**Figure 4C**). Of the 392 such interactions, 53% are already listed as known ‘associations’ between homologous proteins in the STRING database (Szklarczyk et al., 2021) for *M. genitalium* (***Methods*, Table S3**). And, of the 167 predicted detectable interactions excluding aaRS-tRNAs (**Figure 4B, Table S2**), 69% are listed in STRING.

## Discussion

We used dynamic simulation to interrogate how ultra-weak PPIs (UW-PPIs) could influence ensemble protein behavior in prokaryotic cytoplasm. Explicit representation of deterministic attractions, entropic exclusion, and Brownian motion enabled our computational model to capture the transient spatio-temporal dynamics underlying UW-PPIs. Moreover, the added spatial and temporal resolution of our model revealed heterogeneous mesoscale dynamics, namely the formation of transient clusters that arise from UW-PPIs, dissociate over time due to thermal fluctuations, and re-form into new clusters (**Figure 2A**). Such phenomena can facilitate combinatoric sampling of multi-molecular interactions and cannot be predicted by mass-action kinetics alone.

We first established the baseline connection between entropic exclusion (i.e., in the total absence of attractive forces) and binding affinity, which produced a quantifiable apparent affinity of *K*_*d,app*_ ∼20 mM (**Figure 2B**). This underlying affinity simply reflects the tendency of Brownian motion and crowding to facilitate protein encounters, which are tracked explicitly in our spatially-resolved simulations. We purposefully used the word encounter to describe such events where proteins are in sufficiently close proximity to enable chemical reactions (Idziaszek et al., 2015; Salam, 2010) yet there are no interaction-specific forces to favor one encounter over another.

We found that attractive UW-PPIs below current experimental detection limits should drive transient protein co-localization and micro-condensate formation while avoiding space-spanning gelation (**Figure 2B, S2, S3**). By tracking positions of individual proteins, we demonstrated that UW-PPIs can both increase the likelihood and accelerate sampling of protein associations via the formation of transient clusters, which last an order of magnitude longer than in the zero-attraction limit yet can still dissociate and allow proteins to search for new partners (**Figure 3, S7**). We found that ultra-weak dimers remain intact long enough for elementary chemical reactions (Gruebele and Zewail, 1990) and protein conformational fluctuations (Bredenbeck et al., 2003) to occur, whereas larger ultra-weak clusters are durable enough to facilitate the fastest known enzyme-catalyzed reactions (Smejkal and Kakumanu, 2019) (**Figure 5**, left). Cytoplasmic transport times – i.e., how long it takes for a protein to diffuse and encounter its nearest neighbor or desired binding partner – are also comparable to the duration of ultra-weak clusters. For example, an EF-Tu::GFP::tRNA ternary complex (with a radius of 5.9 nm (Maheshwari et al., 2022) and *in vivo* diffusivity of 2.2 *μ*m^2^/s (Mustafi and Weisshaar, 2018)) will take ∼2 *μ*s to diffuse the 5.5 nm to its nearest ribosome, or ∼76 ns to diffuse the 1.0 nm to its nearest macromolecular neighbor at a cytoplasm volume fraction of 30% (Maheshwari et al., 2022). This ultra-weak, transient clustering regime also operates below the characteristic timescales of whole-cell and macroscopic phase separation processes (Asthagiri and Lauffenburger, 2003; Molliex et al., 2015; Ramm et al., 2022; Tang et al., 2021; Zia et al., 2014) (**Figure 5**, right), which are readily measured *in vivo*.

In extending our analysis to the entire cytoplasmic proteome of *M. genitalium*, we found that only 13% of the isotropic electrostatic attractions predicted by our method would be strong enough to be detected by current *in vivo* methods (**Figure 4C**). Of these predicted detectable interactions, 50-70% are already listed as known ‘associations’ between homologous proteins in the STRING database (Szklarczyk et al., 2021) for *M. genitalium*. Our method might find use in helping to predict electrostatic PPIs for any organism with available proteomic charge data, with an accuracy similar to other computational methods such as AlphaFold-Multimer (Evans et al., 2022) and abstract mining (He et al., 2009; Papanikolaou et al., 2015), both of which have ∼70% accuracy. Unlike abstract mining, our method does not rely on prior literature and should thus be better suited for identifying as-yet unknown interactions. Moreover, by refining the interaction potential, our technique could be extended to model additional types of PPIs including hydrophobic and van der Waals interactions. Further work might also include direct, systematic comparison between computation and experiment across a full range of binding affinities, especially given the current lack of direct PPI measurements for *M. genitalium* proteins.

What can we say about the 87% of electrostatic PPIs in the *M. genitalium* cytoplasm that we predict exist but cannot yet be readily measured via experiment? To better understand possible biological implications, we focused on the 14% of such cytoplasmic PPIs that cause proteins to experience a binding-dominant regime, yet their interaction is just below the current detection threshold (**Figure 4**, binding-dominant UW-PPI region). In this category, we identified specific pairs of interacting proteins that are involved with related biological functions (**Table S3**), including pairs of chaperones, translation-associated proteins, and consecutive metabolic enzymes. We speculate that transient co-localization driven by these UW-PPIs could increase the likelihood and duration of associations while speeding up the diffusive search process (**Figure 3A**) between proteins involved in adjacent biological processes. These transient associations, for example, could facilitate more rapid exchange of protein substrates between the DnaK/DnaJ/GrpE and GroEL/ES chaperones or pyruvate between pyruvate kinase and pyruvate dehydrogenase complex, as has been hypothesized in the case of heme transfer between hemoproteins in the cell wall of *Staphylococcus aureus* (Villareal et al., 2011). Our results should motivate further computational and experimental study to directly test the impact of such interactions on cellular growth and fitness. More generally, the prevalence of binding-dominant, ultra-weak PPIs in a minimal cell such as *M. genitalium* suggests that they could be conserved even among the most basic of life-essential genes and fundamental biological processes.

Our results also suggest that the protein interactome could be restructured by the addition of just one or a few molecules. For example, we found that the presence of the ribosome, its 50S and 30S subunits, and aaRS-tRNA can shift the overall protein interaction distribution to higher attraction strengths and increase the likelihood of binding-dominant UW-PPIs by ∼7-fold. These results both underscore the centrality of protein synthesis machinery in prokaryotic cytoplasm and support the assertion that ribosome surface charge is a primary regulator of protein dynamics *in vivo* (Schavemaker et al., 2017). Given the growth-limiting nature of protein synthesis in prokaryotes (Belliveau et al., 2021; Maheshwari et al., 2022), any PPI that increases the speed of translation is likely to have a fitness benefit. As such, we speculate that improved colloidal-scale transport of translation-associated proteins and complexes driven by UW-PPIs could broadly support efficient protein synthesis and overall cellular growth. It is also interesting to imagine how a single new protein could appear within the evolutionary history of a proteome, or be added purposefully, and cause significant changes to the overall PPI distribution.

In addition to proteome composition and structure, our results also suggest that UW-PPIs could be readily tuned by changes in the physical environment within cells, a phenomenon that has also been observed for *in vivo* liquid-liquid phase separation (Banani et al., 2017). For example, we found that small changes in the microscopic attraction strength between proteins, from zero to 2.5 times the strength of thermal fluctuations, can drive order-of-magnitude changes in apparent binding affinity (**Figure 2B**, *V*_0_/*k*_*B*_*T* ≤ 2.5). Changes in attraction strength of this magnitude could be caused by relatively common physiological changes in salt concentrations (100-500 mM (Spitzer and Poolman, 2009; Szatmári et al., 2020)) or pH (4-7 (Kusova et al., 2021)). Temperature and viscosity could further tune UW-PPIs by slowing down or speeding up diffusive associations, suggesting another potential function of “viscoadaptation” – the homeostatic regulation of viscosity by intracellular trehalose and glycogen (Garcia-Seisdedos et al., 2020; Persson et al., 2020). Thus, UW-PPIs might be uniquely sensitive to modest changes in intracellular conditions relative to strong PPIs, which tend to be regulated explicitly and specifically via gene expression levels or timing, post-translational modifications, or sub-cellular location (Nooren and Thornton, 2003).

More broadly, our work underscores the utility of colloidal physics models for studying protein behavior at length- and timescales currently inaccessible by *in vivo* experiments and all-atom simulations (Maheshwari et al., 2019). In the case of ultra-weak PPIs, such spatio-temporal resolution is essential for capturing and connecting transient associations to ensemble properties and biological functions. Further, our model provides a basis for investigating how protein behavior varies generally with colloidal-scale molecular and cellular properties. For example, confinement (Aponte-Rivera and Zia, 2022; Aponte-Rivera et al., 2018; Gonzalez et al., 2021) (e.g., by a cell membrane) and size polydispersity (Asakura and Oosawa, 1954; Gonzalez et al., 2021; Maheshwari et al., 2022) can induce structural correlations between molecules, which we might expect to augment co-localization driven by UW-PPIs.

We encourage further work at the intersection of colloidal physics and cellular biology, which we expect will reveal new biological questions for future experimental study. We envision that colloidal-scale models of cellular proteomes, like those conducted here, will be used to scaleably identify putative PPIs for further investigation, especially as single-molecule and *in vivo* PPI techniques (Danielsson et al., 2019) continue to advance. Moreover, such models could provide a framework for interpreting in-cell images from cryogenic electron microscopy (Pilhofer et al., 2010) or tomography (Turk and Baumeister, 2020) by inferring details of molecular interactions from the degree and duration of co-localization.

## Methods

### I. Computational model

In this study, we constructed a computational model suspension of monodisperse proteins using the molecular dynamics package LAMMPS (Thompson et al., 2022). Each protein was represented explicitly as a nearly-hard colloidal sphere of diameter *σ* (or radius *a*_*p*_) and immersed in an implicit Newtonian solvent of viscosity *η* and density *ρ* in a periodically replicated domain containing 6912 proteins. Here, we set the protein diameter to be 5.2 nm, representative of the average cytoplasmic protein size in *M. genitalium* (Dutagaci et al., 2021). The solvent viscosity and density were chosen to reflect that of water at 37°C, i.e. 0.6913 mPa/s and 0.99336 g/cm^3^, respectively. We chose a total protein volume fraction *ϕ* = 30%, where 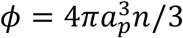 and *n* is the number density of proteins. This volume fraction represents *in vivo* crowding in prokaryotic cytoplasm at moderate growth rate (Maheshwari et al., 2022) Because of the small characteristic size of proteins, the Reynolds number, *Re* = *ρUa*_*p*_/*η*, is vanishingly small and thus fluid motion obeys the Stokes’ equations. Here, *U* is the characteristic protein velocity.

Protein-protein interactions in our model included attractive and entropic components. We represented screened electrostatic attractions between proteins via a Debye-Hückel potential (Debye and Hückel, 1923; Dutagaci et al., 2021). Entropic exclusion (finite size) was enforced via a repulsive Morse potential, which we have previously demonstrated recovers hard-sphere behavior (Zia et al., 2014). The distance-dependent, pair-wise interaction between proteins *i* and (illustration, **Figure S1A**) is

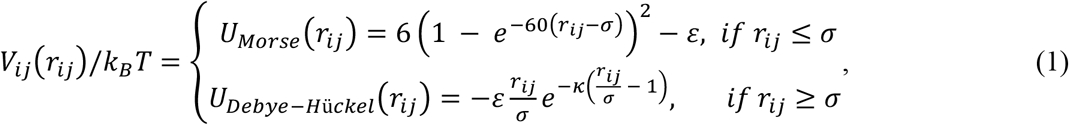

where *r*_*ij*_ is the magnitude of the unit vector 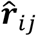 pointing between the center of protein *j* and the center of protein *i, ϵ* = *V*_*0*_*/k*_*B*_*T* is the strength of the electrostatic attraction at contact (in this work, 0 ≤ *ϵ* ≤ 6, see **Table S3**), and *κ*^−1^ is the inverse Debye length that sets the range of the screened electrostatic attraction. We chose *κ* = 7.8 Å, corresponding to the screening length in a ∼150 mM monovalent electrolyte solution, representative of prokaryotic cytoplasm (Record et al., 1998; Spitzer and Poolman, 2009).

Protein motion is governed by the Langevin equation, a stochastic force balance,

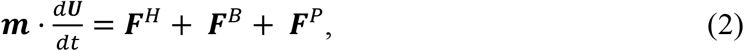

where ***m*** is the mass tensor, ***U*** is the protein velocity, and ***F***^*H*^, ***F***^*B*^, and ***F***^*P*^, are the hydrodynamic drag, Brownian, and interparticle forces, respectively. The hydrodynamic drag on each protein *i* is given by Stokes’ drag law,

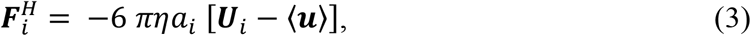

where ⟨***u***⟩ is the bulk fluid velocity which is zero for a quiescent, intracellular suspension. The stochastic Brownian force arising from thermal fluctuations obeys Gaussian statistics:

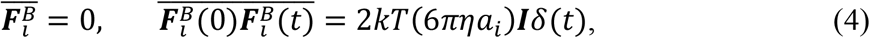

where the overbar denotes an average over times long compared to the solvent timescale, and the Dirac delta distribution *δ*(*t*) reflects the instantaneous correlation of the solvent-molecule impacts that produce Brownian motion. The interparticle force is calculated as a pair-wise sum over all nearby proteins,

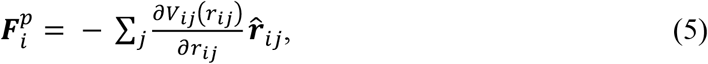

where *V*_*ij*_ is the interparticle potential calculated for each pair according to equation (1).

The Langevin equation (2) can be solved to produce trajectories for all proteins in simulation as they interact with each other. In LAMMPS, protein velocities were integrated forward in time via the velocity-Verlet method (Allen and Tildesley, 2017) and a Langevin thermostat (Brünger et al., 1984). The inertia-less physics of the Stokes-flow regime were recovered by setting a small damping factor d = 0.01 and discretization timestep Δ*t* = 2 ps (Zia et al., 2014). Simulations were evolved for 55 μs, long enough for the densely-packed suspension to be well mixed. The simulations output a set of highly-resolved protein trajectories: position versus time for all proteins in the simulation, written to file at each 2 ps timestep.

Validation of our method is obtained by simulating passive Brownian diffusion: the explicit mean-square displacement of each protein was calculated and the resulting *ab initio* long-time self diffusion coefficient 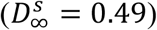 was compared to prior theoretical 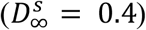 (Brady, 1994) and computational measurements 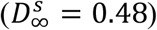 (Cichocki and Hinsen, 1992). We emphasize that diffusion in our simulations emerges from first-principles physics: thermal solvent fluctuations and entropic exclusion produce a physical random walk through space, in contrast to kinetics-only models which implement an effective diffusivity as a parameter.

### II. Calculation of apparent binding affinity, K_d,app_

We calculated apparent equilibrium dissociation constants, *K*_*d,app*_, termed ‘apparent’ to follow previous definitions for concentrated systems (Paramanathan et al., 2014; Sarkar, 2020). In our dynamic simulations, we computed the *K*_*d,app*_ of protein-protein interactions (**Figure 2**) via the relation (Duffey, 2000):

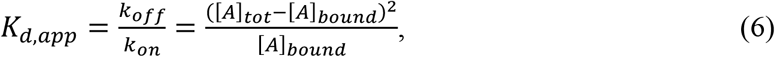

where *k*_*off*_ and *k*_*on*_ are the dissociation and association rate constants, [*A*]_*tot*_ is the total protein concentration, and [*A*]_*bound*_ is the concentration of proteins found in binding events. In this work, we defined two proteins as ‘bound’ when their surfaces are separated by less than 0.5 Å (i.e., 0.01*σ*, where *σ* is the protein diameter). This surface-to-surface separation criterion, *r*_*cut*_, corresponds to the atomic length scale at which chemical reactions occur (Idziaszek et al., 2015; Jachymski et al., 2013) and where quantum-mechanical effects will dominate longer-ranged, coarse-grained interaction potentials (Bian et al., 2021; Roth et al., 1996; Salam, 2010). To establish the sensitivity of binding events to the choice of separation criterion, *r*_*cut*_, we conducted sensitivity analysis of *K*_*d,app*_ and all other measured data (see **Figures S1, S2, S4, S8**) and found that our qualitative results do not change.

We equilibrated the system for 54 μs to allow Brownian motion and binding events to reach steady state. In the final 300 ns of that simulation, we extracted binding data to calculate binding affinities *K*_*d,app*_ via equation (6).

### III. Calculation of second virial coefficients

We calculated the second virial coefficient for an isotropic interaction potential *V*(*r*) as

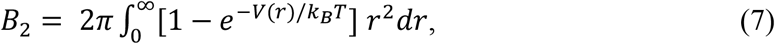

where *r* is the center-to-center separation between proteins. The second virial coefficient *B*_2_ is a thermodynamic descriptor of the deviation of pair interactions from ideal gas behavior (i.e. a system of non-interacting point particles, with *B*_2_ = 0). Normalizing this coefficient on the hard-sphere value 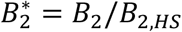, communizes the effect of finite particle size, explicitly revealing the tendency of other interactions to cause attraction or repulsion. This normalization, 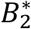, is called the reduced second virial coefficient and predicts phase behavior independent of the detailed shape of *V*(*r*), as long as the potential is short ranged (Noro and Frenkel, 2000). We performed numerical integration of equation (7) for both parts of the potential in equation (1) for 0 ≤ *V*_0_/*k*_*B*_*T* ≤ 6 using the SciPy library in Python.

### IV. Quantification of protein binding dynamics

We quantified the dynamics of individual protein associations by tracking the time proteins spend bound to one another (‘binding events’) and searching for new partners (‘search events’). As described in Methods Section III, we defined a protein binding event to occur when protein surfaces come within *r*_*cut*_ = 0.5Å of hard-sphere contact. We tracked the state (either ‘bound’ or ‘unbound,’ **Figure S5A**) of every protein at each 2 ps timestep, over the full analysis window of 300 ns. At each timestep, a particle was either bound or unbound; when its status changed from bound to unbound or vice-versa, the duration of the departed state was recorded. The durations of these bound and unbound events were recorded for every particle in the system. We calculated the probability distribution of both types of event durations and averaged over the protein ensemble, normalizing so that the total event probability sums to one (**Figure 3, lower**). We chose to focus on data for *t* >200ps in order to monitor diffusive rather than ballistic dynamics, i.e. at timescales long compared to the particle momentum relaxation timescale (3 ps for a protein in water at 37°C). We confirmed diffusive behavior by showing that mean squared displacement grows linearly with time (**Figure S6**). We conducted a sensitivity analysis of our results to the choice of separation criteria, *r*_*cut*_, provided in the Supplementary Information (**Figures S8**).

We calculate average cluster dissociation times (**Figure S5B**) by identifying all clusters comprising three or more proteins at the beginning of the analysis period. Two particles are said to be in the same cluster if their surfaces are separated by a distance less than *r*_*cut*_ = 0.01*σ*. We tracked protein positions at each timestep and calculate the length of time required for all proteins in each initial cluster to separate (**Figure S7B**), i.e., for the cluster to fully dissociate.

### V. Extension to *M. genitalium* proteome

Effective charges of cytoplasmic proteins in *Mycoplasma genitalium* were obtained from (Dutagaci et al., 2021). The strength of the electrostatic interaction between a pair of proteins (i,j) is defined as:

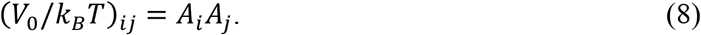

The form of the modified strength parameter *A*_*i*_ was parameterized against atomistic simulations in the previous work by Dutagaci and colleagues as:

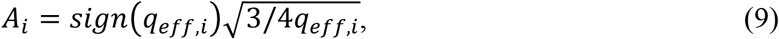

where *q*_*eff,i*_ is the total effective charge of protein *i*. A list of *M. genitalium* cytoplasmic proteins, their associated charges *q*_*eff,i*_, strength parameters *A*_*i*_, and all calculated (*V*_0_/*k*_*B*_*T*)_*ij*_ values are provided in **Supplementary File 1**. We focused on electrostatic attractions in this work and thus limited our analysis to pair-wise interactions with (*V*_0_/*k*_*B*_*T*)_*ij*_ <0, i.e. protein pairs with opposite charges. Knowing the set of (*V*_0_/*k*_*B*_*T*)_*ij*_ for all cytoplasmic electrostatic PPIs, we calculated the probability distribution of attraction strengths, *P*((*V*_0_/*k*_*B*_*T*)_*ij*_ ∣ *i* = *k*), experienced by each protein *k* (each row in **Figure 4, lower**), as well as the overall distribution across all pairs *P*((*V*_0_/*k*_*B*_*T*)_*ij*_).

### VI. Comparison to homologous protein associations in STRING

As another stress test of our model, we compared each PPI we predict to be experimentally detectable (**Tables S1-S3**) against known protein-protein associations in the STRING database (Szklarczyk et al., 2021) for *Mycoplasma genitalium* (NCBI ID: 243273). We defined two proteins as interacting if their STRING interaction score was greater than 0.4 (medium confidence) and included all possible interaction sources: textmining, experiments, databases, co-expression, neighborhood, gene fusion, and co-occurrence measurements. We defined a protein-ribosome interaction to exist if a protein is associated with one or more ribosomal protein in STRING. Because genomic context predictions are currently the only direct evidence of protein associations in *M. genitalium*, we also include associations for putative homologs in other organisms.

## Acknowledgements

We thank Emma Gonzalez and Theo Yang for valuable discussions regarding macromolecular dynamics, suspension mechanics, and experimental quantification of protein-protein interactions. We thank Sasha Levy for providing helpful comments on the manuscript. This work was supported by National Science Foundation Graduate Research Fellowships under Grant No. – 1656518 for J.L.H. and A.M.S., a National Institutes of Health T32 Training Grant No. GM007365 for A.J.M., as well as a Stanford Graduate Fellowship for A.M.S. The computing for this project was performed on the Sherlock cluster. We thank Stanford University and the Stanford Research Computing Center for providing computational resources and support that contributed to these research results.

## Supplemental Information

### Note S1. Choice of separation criterion for protein binding

In this work, we defined a protein as bound when its surface comes within 0.5 Å or 0.01*σ* of hard-sphere contact with another protein, where *σ* is the protein diameter, corresponding to the atomic length scale at which chemical reactions occur and where atomic details will dominate colloidal-scale physical attractions (see ***Methods***). At this separation, electrostatically attractive proteins experience a strong deterministic force relative to the strength of thermal fluctuations (**Figure S1A**) that will tend to pull pairs towards contact. The chosen binding criterion also corresponds closely with the nearest-neighbor entropic peak in the radial distribution function g(r/*σ*), shown in **Figure S1B**. The radial distribution function describes the pair-level microstructure due to correlations in particle positions arising from the finite-size nature of macromolecules. Defining a binding criterion close to the nearest-neighbor peak provides a first-order effect of entropic exclusion, whereas a larger cutoff value would artificially compound entropic effects of dense (volume fraction *ϕ* = 30%) suspensions. Our chosen binding criterion also corresponds to the average protein-protein bond length in the highest affinity case studied here (**Figure S2**) and is thus a fair representative definition of a protein binding event. We conducted a sensitivity analysis of our conclusions to the choice of binding criteria closer to hard-sphere contact, the results of which are shown in **Figures S4** and **S8**.

### Note S2. Coordination number distributions

To quantify the microstructural effects of ultra-weak PPIs, we calculated the distribution of coordination numbers experienced by proteins in dense colloidal suspensions (**Figure S3**). Two particles are coordinated if they are bound to one another, i.e., their surfaces are found within a separation distance of 0.01*σ*. In the hard-sphere, zero-attraction case (*V*_0_/*k*_*B*_*T* = 0), proteins tend not to be coordinated with any neighbors, with only 17% of proteins having a coordination number greater than zero. A PPI at the ultra-weak threshold (*V*_0_/*k*_*B*_*T* = 2.5) drives moderate structuring, with nearly 50% of proteins experiencing coordination numbers between 1 and 5. Moderate-affinity PPIs (*V*_0_/*k*_*B*_*T* = 6) drive formation of dense, durable networks where proteins are coordinated with many neighbors, up to ten each. In the moderate-affinity case, less than 5% remain uncoordinated, compared with more than 80% in the hard-sphere, zero-attraction case.

### Note S3. Sensitivity of structural results to the binding separation criterion

We examined the sensitivity of our conclusions to variations in the binding separation criterion, comparing results using a value of *r*_*cut*_ = 0.01*σ*, as in the reported data, to smaller values of *r*_*cut*_ = 0.008*σ* and 0.005*σ* (**Figure S4**). We calculated the apparent equilibrium dissociation constant via:

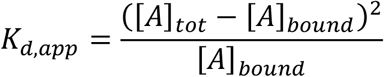

where [*A*]_*tot*_ is the total protein concentration and [*A*]_*bound*_ is the concentration of bound protein. Thus, [*A*]_*bound*_ is sensitive to the definition of a binding event, set by the separation distance *r*_*cut*_. We found that *K*_*d,app*_ is only minimally affected by the definition of a binding event (*r*_*cut*_ less than 0.01*σ*) in systems with ultra-weak binding affinities, our regime of interest. The calculated binding affinity was more affected by the binding separation criterion in systems with weak to moderate attractions (*V*_0_/*k*_*B*_*T* greater than 3), beyond the ultra-weak regime. If interactions are isotropic, the equilibrium assumption begins to break down in this limit with the formation of arrested gel networks. Additional analysis is needed in this regime to match computational models to non-equilibrium experimental systems (Ryu et al., 2022).

Ultimately, our overall claims are insensitive to the choice of stricter binding separation criteria (*r*_*cut*_ less than 0.01*σ*). For all three studied criteria, an apparent binding affinity of *K*_*d,app*_ ∼20 *mM* arises in purely repulsive systems, driven by the entropic tendency of particles in dense suspensions to occasionally sample positions near one. In addition, weak attractions up to 2.5 times the strength of thermal fluctuations *k*_*B*_*T* drive an approximately order-of-magnitude change in the apparent binding affinity experienced by proteins, for all three separation criteria.

### Note S4. Cluster size distributions and validation of *K*_*d,app*_

Since our calculation of *K*_*d,app*_ in the ultra-weak regime cannot be validated against experiments, we turned to a recent theoretical relationship between the mean protein cluster size and binding affinity developed by von Bülow and colleagues for equilibrium systems (von Bülow et al., 2019). Their result, 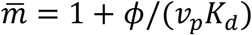, where 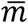 is the average cluster size, *ϕ* is the overall volume fraction, and *v*_*p*_ is the protein volume, was shown to hold up to moderate protein concentrations of 100 mg/ml via comparison against all-atom simulations. Though our colloidal simulations at volume fraction *ϕ* = 30% (∼300 mg/ml) are more concentrated, such a comparison is a useful reference point to qualitatively validate our experimentally inaccessible results.

Here, we defined a cluster as a set of particles whose surfaces are found within the specified binding cutoff distance of at least one other particle in the cluster. We chose a cutoff distance of 0.01*σ*, where *σ* is the particle diameter, consistent with our earlier results (**Figures 2-4**), and calculated the probability distribution of cluster sizes, with average 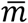 (**Figure S7A**), for systems with a range of ultra-weak binding affinities. We then input these average cluster sizes into the *K*_*d,app*_ relation from von Bulow et al., the results of which agree well with our earlier calculation using the concentration of bound protein (**Figure S7C**).

### Note S5. Sensitivity of dynamic results to the protein binding separation criterion

We examined the sensitivity of our conclusions to variations in the binding separation criterion, comparing results using a value of *r*_*cut*_ = 0.01*σ* (**Figure S8A**), as in the reported data, to smaller values of *r*_*cut*_ = 0.008*σ* (**Figure S8B**) and 0.005*σ* (**Figure S8C**). For each value of *r*_*cut*_, we calculated the distribution of binding and search event durations (left) and overall bound likelihood (right) as a function of PPI binding affinity.

As expected, proteins spend less time in binding events as the separation criteria *r*_*cut*_ becomes stricter (left); protein pairs can only travel shorter distances over shorter timescales. However, our overall claims are insensitive to the choice of stricter binding criteria (*r*_*cut*_ less than 0.01*σ*). In the hard-sphere zero-attraction case, binding events are shorter on average than search events for all binding separation criteria. In the case with affinity at the threshold of the ultra-weak regime, the average search and binding durations remain approximately equal. With moderate-affinity attractions, binding time still dominates search time. With stricter binding criteria, proteins are found to have a decreased likelihood of being found in a binding event (right) – as expected. However, the special intermediate regime that we term ‘binding-dominant’ UW-PPIs, where Da < 1 and *K*_*d,app*_> 1 mM, is still present for all binding criteria.

**Figure S1.**
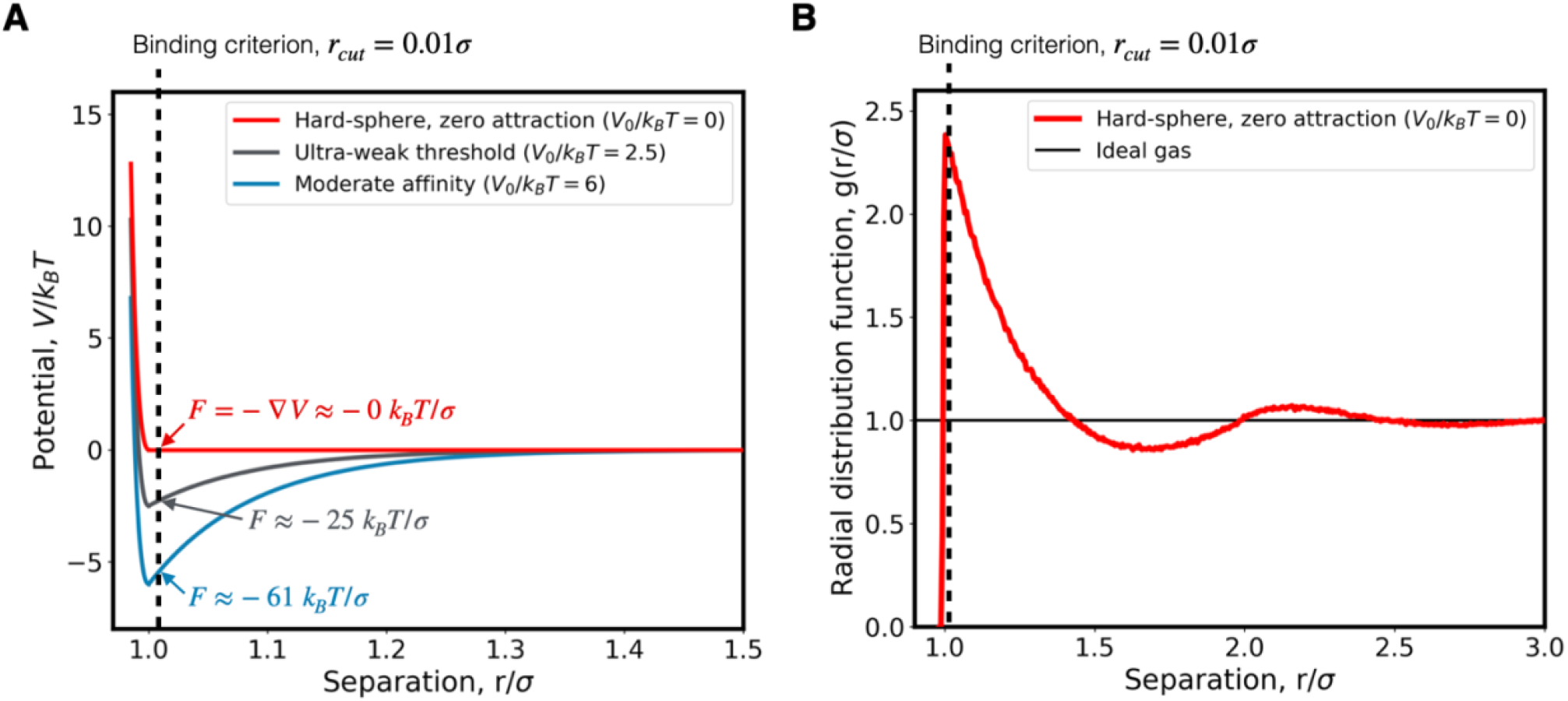
The criterion chosen to specify protein binding is close to the minimum interaction well-depth and protein-protein contact. **(A)** The interaction between proteins in colloidal simulation is represented by an interparticle potential, *V*/*k*_*B*_*T*, that is a function of the interparticle separation distance *r*/*σ*, where *σ* is the protein diameter. Proteins entropically exclude one another (steep region, left), and the attractive binding well deepens as PPI affinity is increased (colors; legend, inset). The chosen binding separation criterion (dashed vertical line) is close to the well-depth that represents protein-protein contact. At this separation, attractive proteins experience strong forces (*F*, marked) that tend to keep a protein pair bound together. **(B)** The probability of finding a protein pair at a given separation (i.e., the radial distribution function, g(*r*/*σ*)) for the simulated hard-sphere, zero-attraction case (red) and an ideal gas of point particles (black). Proteins entropically exclude one another, driving an increased likelihood of finding a protein close to contact (peak at *r*/*σ* = 1) as well as surrounding layers of sequentially lower order (peaks and troughs at higher values of *r*/*σ*). The chosen binding separation criterion (dashed vertical line) is close to the first nearest-neighbor peak.

**Figure S2.**
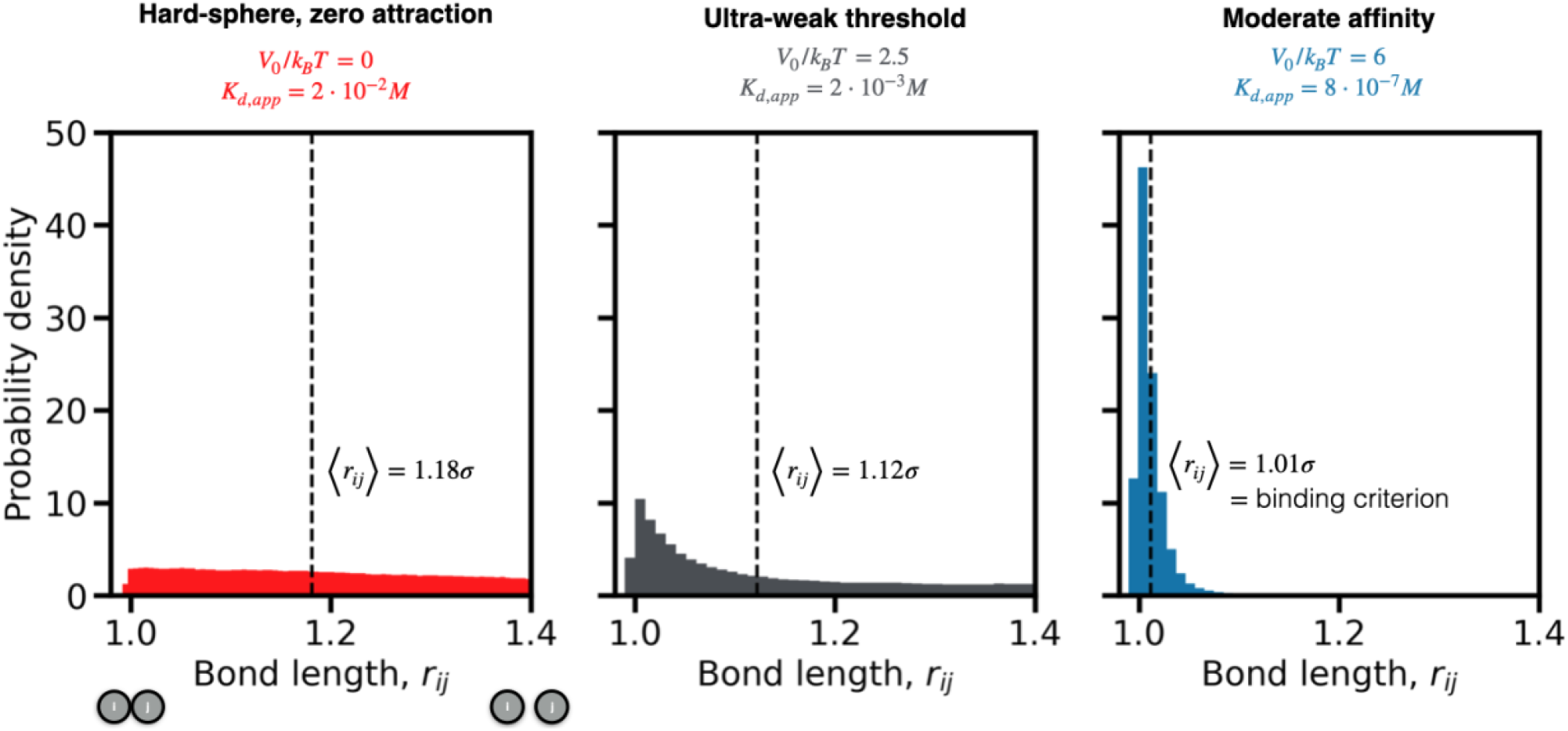
Attractions decrease the average protein-protein ‘bond’ length. The likelihood of finding a protein pair (*i, j*) at a given separation distance *r*_*ij*_ shifts to smaller bond lengths as affinity increases (colors, left to right). The average bond length (dashed line) decreases from 9 Å in the hard-sphere case (red, 1.18*σ*, where *σ* is the protein diameter of 5.2nm), to 6 Å in the ultra-weak threshold case (grey, 1.12*σ*) and 0.5 Å in the moderate affinity case (blue, 1.01*σ*).

**Figure S3.**
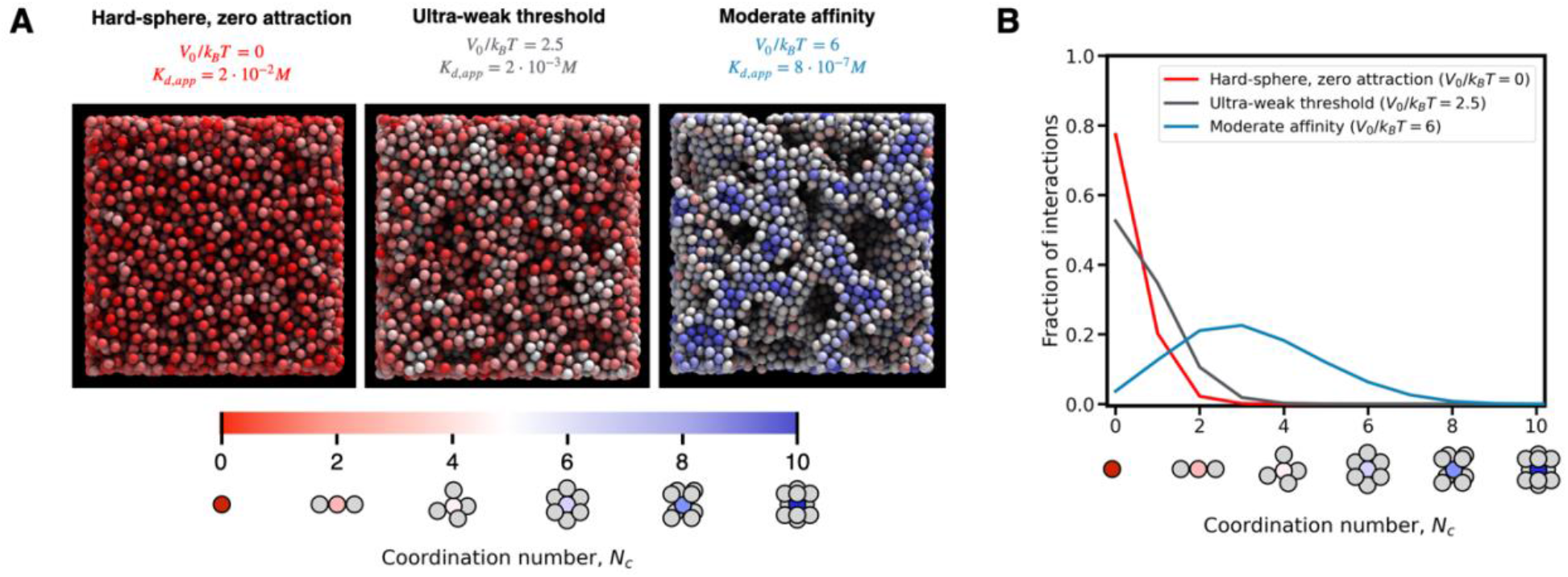
Higher affinity attractions increase the protein coordination number. **(A)** Increasing the strength of attractions (left to right) creates microstructure in simulation snapshots, where proteins are coordinated with increasing number of neighbors (colored; legend, below). **(B)** Ensemble coordination number distributions are heterogeneous, and shift to higher coordination numbers as attraction strength increases.

**Figure S4.**
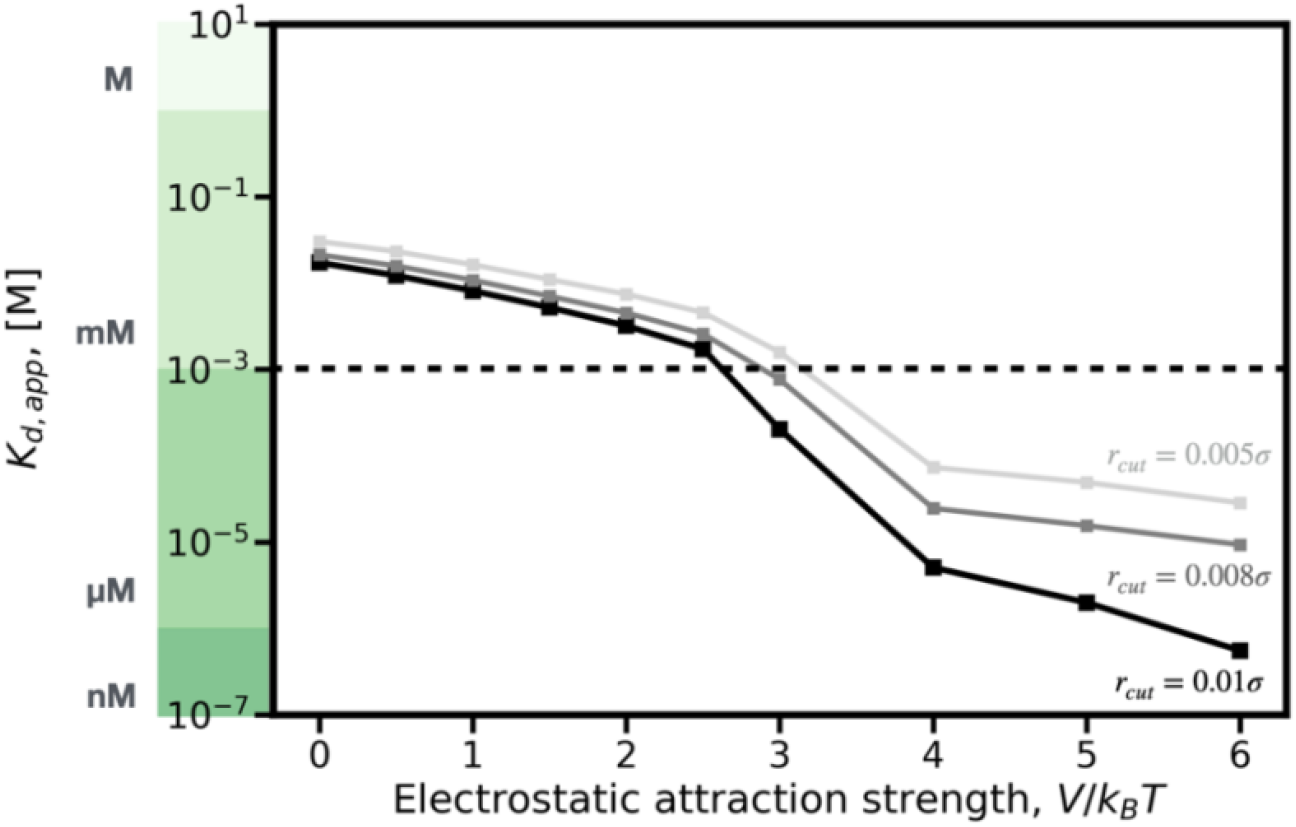
Apparent binding affinity is not qualitatively impacted by the protein binding separation criterion in the ultra-weak regime. Calculation of the apparent equilibrium dissociation constant *K*_*d,app*_ for more strict definitions of a protein binding event (*r*_*cut*_, light to dark grey), compared to the definition used in this work (black).

**Figure S5.**
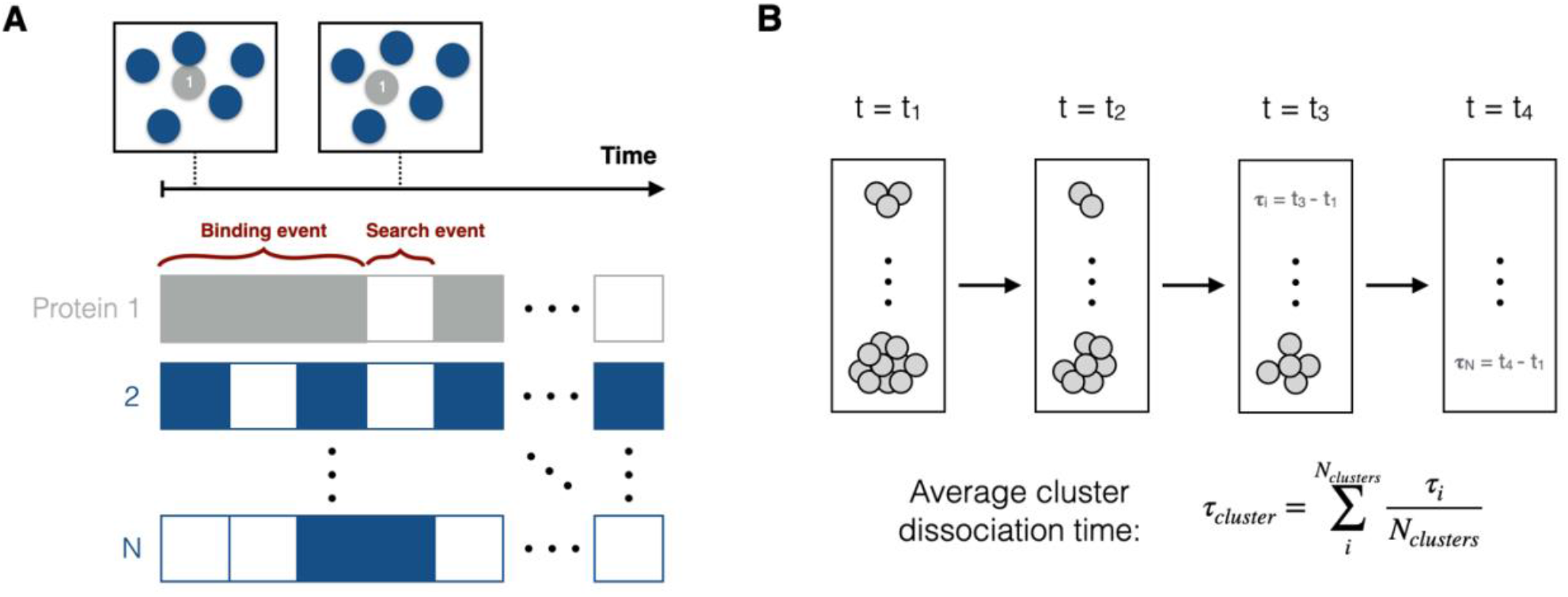
Protein dynamics can be quantified by tracking pair-level binding and search times, as well as meso-scale cluster dissociation times. **(A)** In a suspension (upper), a protein is either bound to another protein or free and searching for a new binding partner. We quantify the duration of ‘binding’ and ‘search’ events for all N proteins in the suspension over time. **(B)** Schematic of the calculation of the average cluster dissociation time. Clusters of particles bound to one another are first identified at t_1_. Two particles are said to be in the same cluster if their surfaces are separated by a distance less than 1% of a particle diameter (*r*_*cut*_ = 0.01*σ*). We then calculate the length of time required for each cluster to dissipate, i.e., for all particles in each cluster at t_1_ to leave said cluster. On average, smaller clusters have faster dissociation times, as expected.

**Figure S6.**
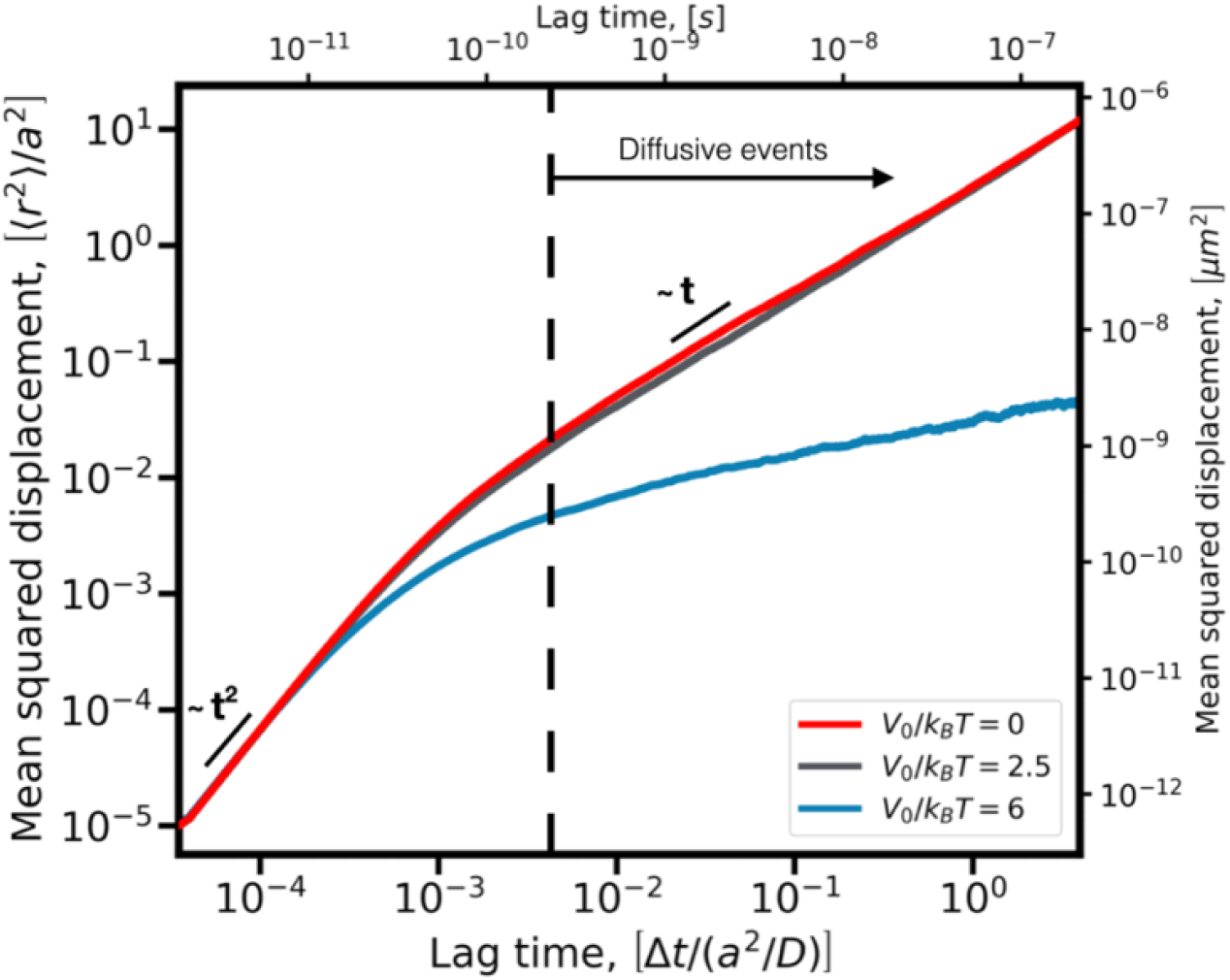
Proteins experience inertial dynamics over short times and diffusive dynamics over longer times. The mean squared displacement (MSD) for proteins in the hard-sphere zero-attraction (red), ultra-weak threshold (grey), and moderate affinity (blue) cases. Particles experience two types of dynamics: inertial behavior at short times (scaling as ∼t^2^), and diffusive behavior at moderate to long times (scaling as ∼t). Analysis of protein-protein binding and search events in this work is limited to timescales greater than 200 ps to accurately capture dynamics in the diffusive regime. Here, *a* is the protein radius and *D* is the protein’s Stokes-Einstein diffusivity defined previously (Methods).

**Figure S7.**
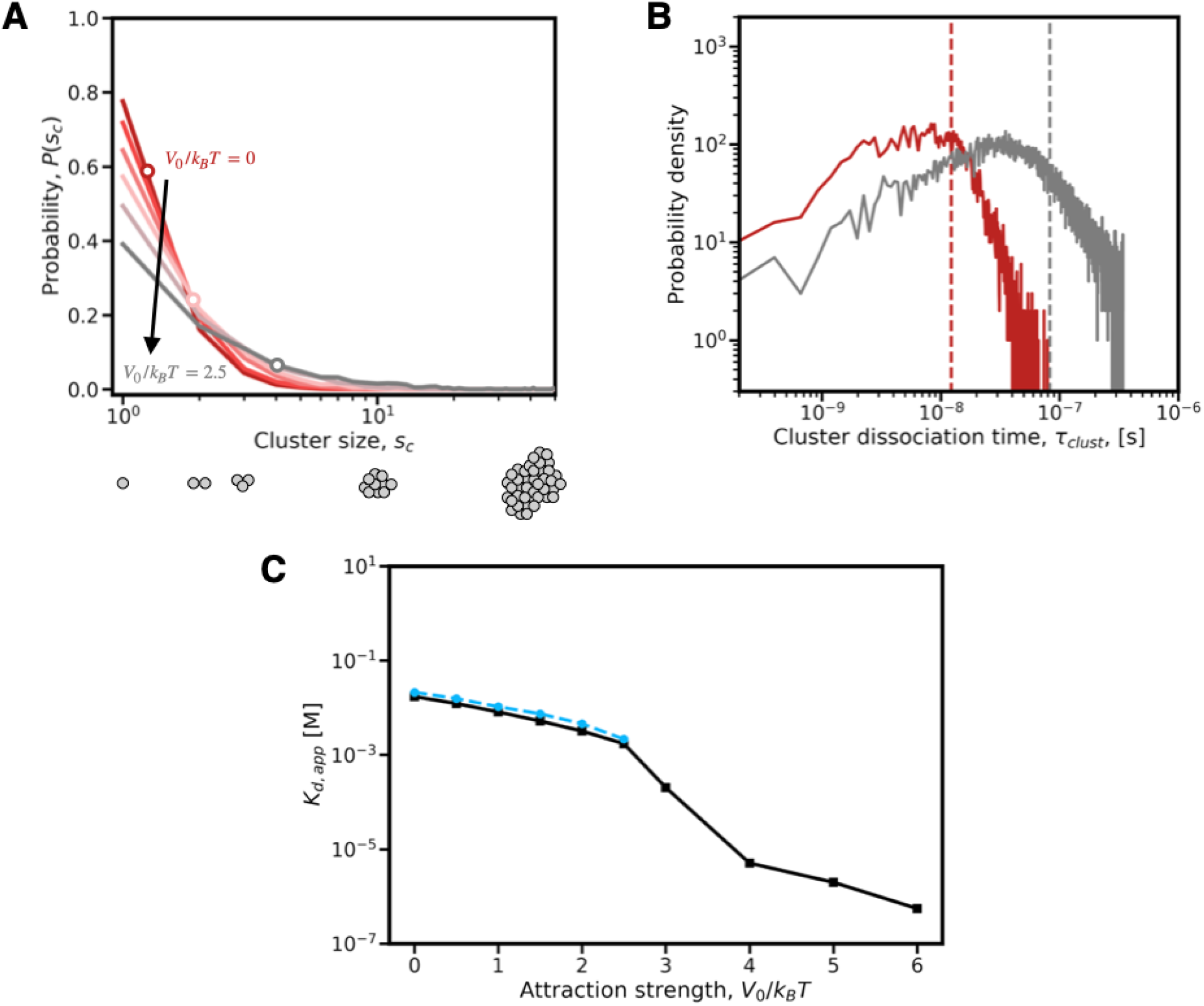
Ultra-weak attractions drive moderate, relatively durable clusters, the size of which can be used to validate the calculation of *K*_*d,app*_. **(A)** The probability distribution of cluster sizes for systems with a range of ultra-weak binding affinities, from the hard-sphere zero attraction case (red, *V*_0_/*k*_*B*_*T* = 0) to the ultra-weak threshold case (grey, *V*_0_/*k*_*B*_*T* = 2.5). The average cluster size (open circles) increases from 1.3 to 4.2 particles over the ultra-weak affinity regime. **(B)** The probability density function of dissociation times for clusters of size 3 or greater in the zero-attraction (red) and ultra-weak threshold (grey) cases. The average cluster dissociation time (vertical dashed line) increases by nearly an order of magnitude over the ultra-weak affinity regime. **(C)** *K*_*d,app*_ as a function of attraction strength, calculated from colloidal simulations via the concentration of bound protein (black, same as in Figure 2), and as a function of average cluster size (blue).

**Figure S8.**
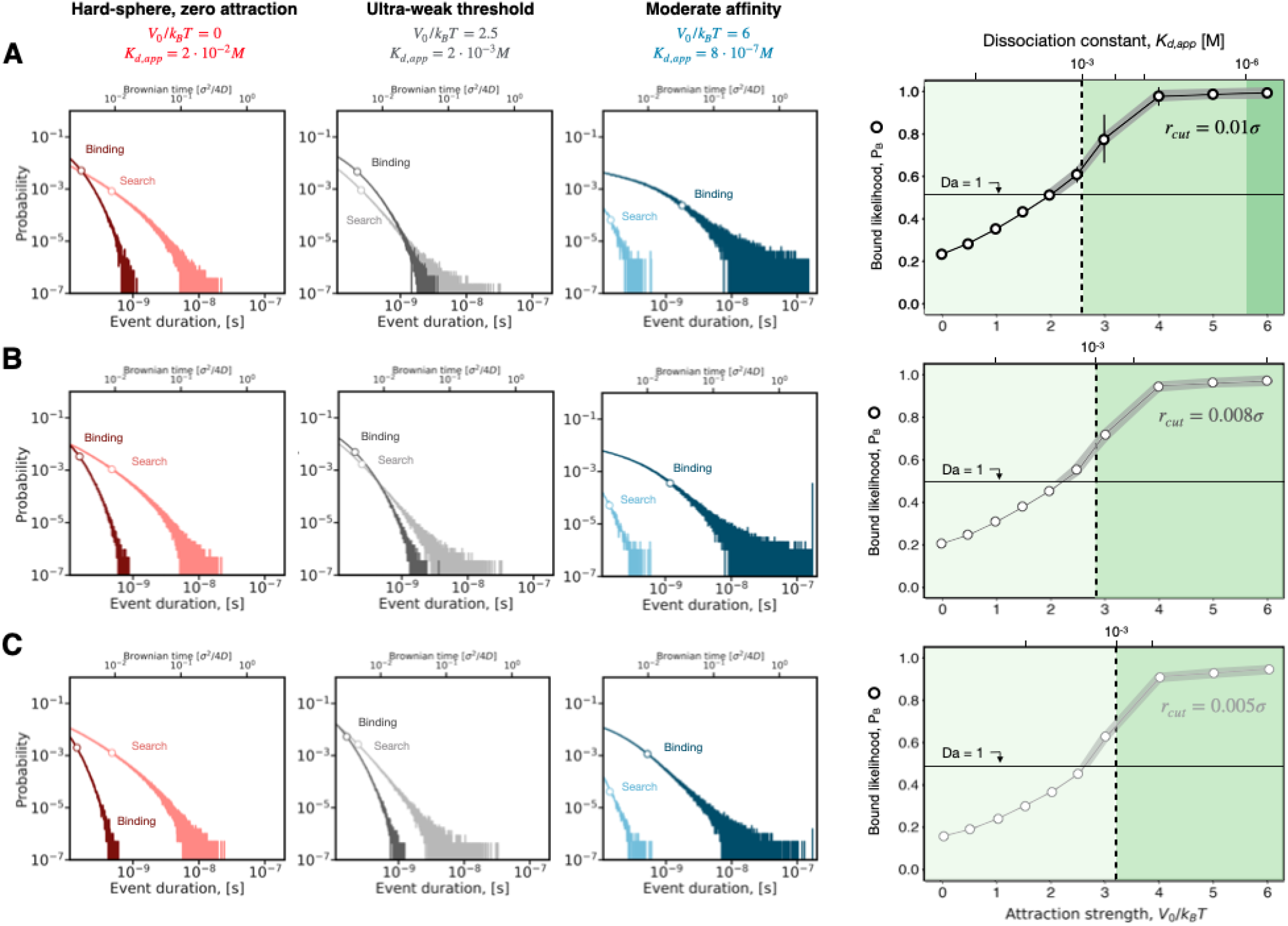
Our overall claims regarding protein binding dynamics are insensitive to the definition of a protein binding event. The distribution of search event durations (left, light colors) and binding event durations (left, dark colors), and the overall likelihood of finding a protein in an binding event (right), for three binding separation criteria **(A)** *r*_*cut*_ = 0.01*σ*, **(B)** *r*_*cut*_ = 0.008*σ*, and **(C)** *r*_*cut*_ = 0.005*σ*. Our claims regarding the relative position of the average search and binding times (left, circles) hold for all binding separation criteria and affinities, as does the ‘binding-dominant UW-PPI region (right; grey shaded region above horizontal line corresponding to Damköhler number Da = 1, and to left of vertical line).

**Figure S9.**
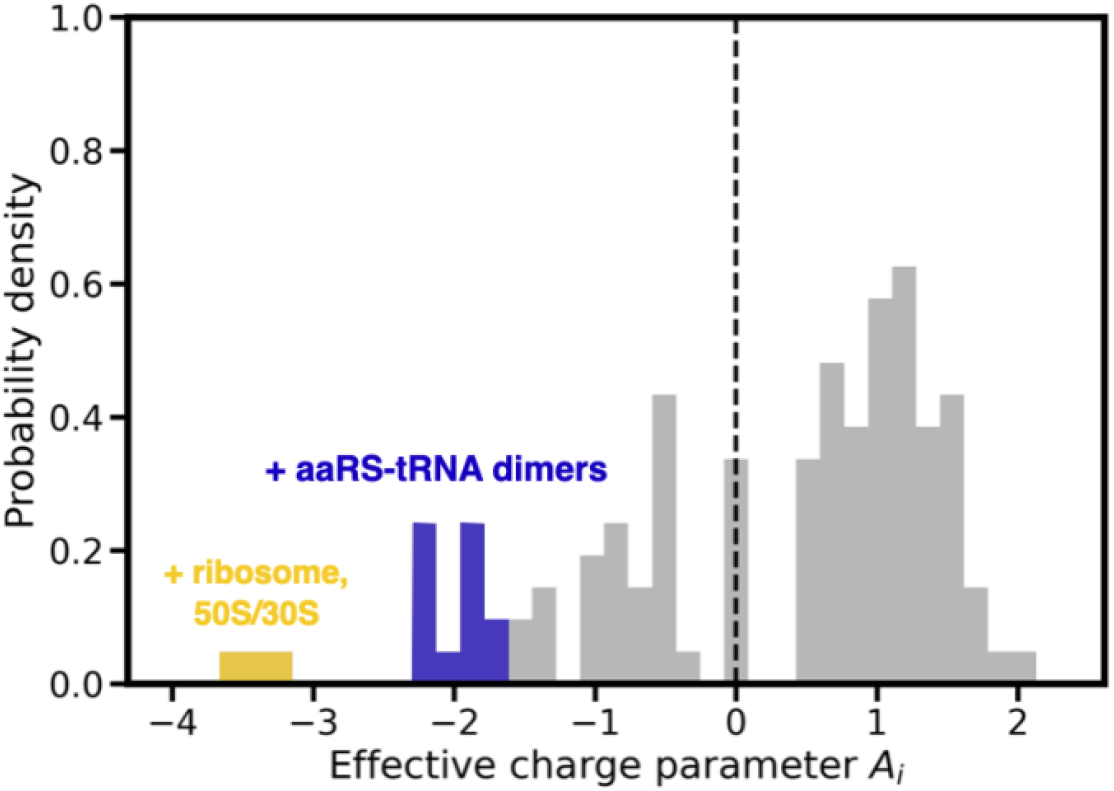
The *M. genitalium* cytoplasmic proteome has a varied charge distribution. The probability density function of the effective charge parameter, *A*_*i*_, for all proteins in the *M. genitalium* cytoplasm, relative to the neutral case (vertical dashed line). See **Methods** for a description of the effective charge parameter and its use in the model.

